# *De novo* apical domain formation inside the *Drosophila* adult midgut epithelium

**DOI:** 10.1101/2021.12.10.472136

**Authors:** Jia Chen, Daniel St Johnston

## Abstract

In the adult *Drosophila* midgut, basal intestinal stem cells give rise to enteroblasts that integrate into the epithelium as they differentiate into enterocytes. Integrating enteroblasts must generate a new apical domain and break through the septate junctions between neighbouring enterocytes, while maintaining barrier function. We observe that enteroblasts form an apical membrane initiation site when they reach the septate junction between the overlying enterocytes. A space appears above the enteroblast as Cadherin clears from its apical surface. New septate junctions then form laterally with neighbouring enterocytes and the AMIS develops into pre-apical compartment with a fully-formed brush border. Finally, the overlying septate junction dissolves and the pre-enterocyte reaches the gut lumen. Enteroblasts therefore form an apical domain before they have a free apical surface. The process of enteroblast integration resembles lumen formation in mammalian epithelial cysts, highlighting the similarities between the fly midgut and mammalian epithelia.

## Introduction

Like the mammalian gut, the *Drosophila* midgut functions to absorb nutrients and acts as a barrier to pathogens(Miguel-Aliaga et al., 2018). The adult midgut consists of a single layer of polarised epithelial cells, with their apical sides facing the gut lumen and basal sides contacting the extracellular matrix (ECM) and muscle layer(Baumann, 2001; Chen and Sayadian, 2018; Shanbhag and Tripathi, 2009). The epithelium is predominantly composed of enterocytes (enterocyte, 90%) and also contains entero-endocrine cells (ee) and their progenitors. The intestinal stem cells and their progeny, the enteroblasts reside on the basal side of the epithelium beneath the enterocytes (Goulas et al., 2012; Micchelli and Perrimon, 2006; Ohlstein and Spradling, 2006). Quiescent enteroblasts are morphologically indistinguishable from ISCs until they are activated to differentiate in response to damage or nutrient-dependent signals (Nászai et al., 2015; O’Brien et al., 2011; Rojas Villa et al., 2019). Over the past 15 years multiple signaling pathways have been found to control ISC proliferation and differentiation, but much less is known about how differentiating enteroblasts insert into the midgut epithelium and polarize to form an apical domain(Antonello et al., 2015; Chen et al., 2016; He et al., 2018; Jiang and Edgar, 2011; Sasaki and Nishimura, 2021; Wang and Hou, 2010; Wu et al., 2021).

The adult midgut epithelium differs from all other *Drosophila* epithelia in several key respects. Firstly, it is a secondary epithelium that is derived from the endoderm, in contrast to most *Drosophila* epithelia which are directly descend from the cellular blastoderm epithelium(Pitsidianaki et al., 2021; Tepass and Hartenstein, 1994; Yarnitzky and Volk, 1995). Secondly, the apical-basal polarity of the epithelium does not require the conserved epithelial polarity factors that polarize all ectodermal and mesodermal epithelia, although most of these factors are expressed in the midgut. Instead, the polarity of the midgut enterocytes depends on the interaction between the integrin adhesion complex and the basement membrane(Chen and Sayadian, 2018). Thirdly, the midgut cells polarise in a basal to apical direction as enteroblasts differentiate into enterocytes as they integrate from the basal side. By contrast, the other well-characterised *Drosophila* epithelia, such as the cellular blastoderm and the follicular epithelium form in an apical to basal direction(Müller, 2018). Finally, the midgut epithelium has an inverted arrangement of lateral junctions to other *Drosophila* epithelia. The occluding, smooth septate junctions form at the top of the lateral domain above typical adherens junctions, containing E-cadherin (Ecad), α-catenin and Armadillo (β-catenin)(Baumann, 2001; Lane and Skaer, 1980; Tepass and Hartenstein, 1994; Tepass et al., 2001).

The smooth septate junctions are composed of the transmembrane proteins Mesh, Snakeskin (Ssk), Tsp2a and Hoka and the scaffolding protein, Coracle (Cora), and provide the barrier to paracellular diffusion (Furuse and Izumi, 2017; Izumi et al., 2021; Izumi et al., 2016; Izumi et al., 2012). The ISCs and early enteroblasts lie below the septate junctions between the enterocytes and only form adherens junctions with neighbouring cells, while mature enterocytes form both adherens junctions and septate junctions, as well as specialised tri-cellular junctions at the vertex between three cells. As an enteroblast differentiates, it therefore needs to develop new septate junctions with the neighbouring enterocytes as it inserts into the epithelium. This also means that the existing septate junctions between neighbouring enterocytes must be broken to allow the new cell to integrate. Furthermore, the barrier function of the epithelium must be maintained during this process to prevent the contents of the gut lumen, such as digestive enzymes and potential pathogens, from leaking into the body.

As enteroblasts integrate into the epithelium and become enterocytes, they form a new apical domain with its characteristic brush border. The formation of an apical domain *de novo* has previously been studied in two other contexts. In the *Xenopus* embryonic epidermis, progenitor cells integrate into the outer layer through a process of a fast radial intercalation, in which they form the apical junction and apical domain at the same time. The apical domain is then specified after the cells have reached the lumen(Sedzinski et al., 2016). A different process occurs during apical lumen formation in MDCK cell cysts in 3D culture. After the first division, the new membrane between the two daughter cells forms a structure called the AMIS (apical membrane initiation site), where apical proteins and fluids are secreted to create the lumen(Bryant et al., 2010). AMIS initiation depends on components of the midbody that forms during cytokinesis and on the formation of tight junctions at the lateral margins of the cell contact region(Blasky et al., 2015; Mangan et al., 2016). The site of apical domain formation in inserting enteroblasts cannot be induced by the midbody, however, since the ISCs divide beneath the epithelial cell layer and the enteroblasts are post-mitotic when they integrate.

Here we characterize the process of enteroblast integration into the midgut epithelium. Our analysis reveals that integrating enteroblasts generate an “apical” domain with a brush border before they reach the gut lumen and have a free apical surface. This pre-apical compartment (PAC) forms below the septate junction between the neighbouring enterocytes and contains all apical markers tested. Differentiating enteroblasts also form new septate junctions with the neighbouring enterocytes below the existing septate junction between these enterocytes. The new septate junctions then move apically as the cell breaks through the existing septate junctions to emerge on the apical surface with a fully-formed apical brush border. By contrast, enteroblasts lacking septate junction components fail to integrate and either form no apical domain or form an internal apical domain.

## Results

### ISCs/early enteroblasts are polarised and reside under tri-cellular junctions

To analyse the steps in enteroblast integration, we first characterized the polarity and arrangement of junctions in quiescent enteroblasts and ISCs, which lie on the basal side of the epithelium. At steady state, ISCs, quiescent enteroblasts (>50%) and entero-endocrine cells are diploid (2c/4c)(Rojas Villa *et al*., 2019; Xiang et al., 2017). ISCs and early enteroblasts can therefore be identified by their low nuclear volume and lack of expression of Prospero, which specifically labels entero-endocrine cells (ees)(Micchelli and Perrimon, 2006; Ohlstein and Spradling, 2006). Both cell-types are already polarized, as they localize the apical polarity factor Par-6 in a crescent at the apex of the cell (Figure 1A, 1B and S1C). Par-6 is not required for enterocyte polarity, but nonetheless provides a useful marker for the early polarization(Chen and Sayadian, 2018). The actin cytoskeleton is not polarised at this stage, however, and adherens junctions with the neighbouring enterocytes localize uniformly around the cell-cell contacts (Figure 1D, S1A and S1D).

**Figure 1.**
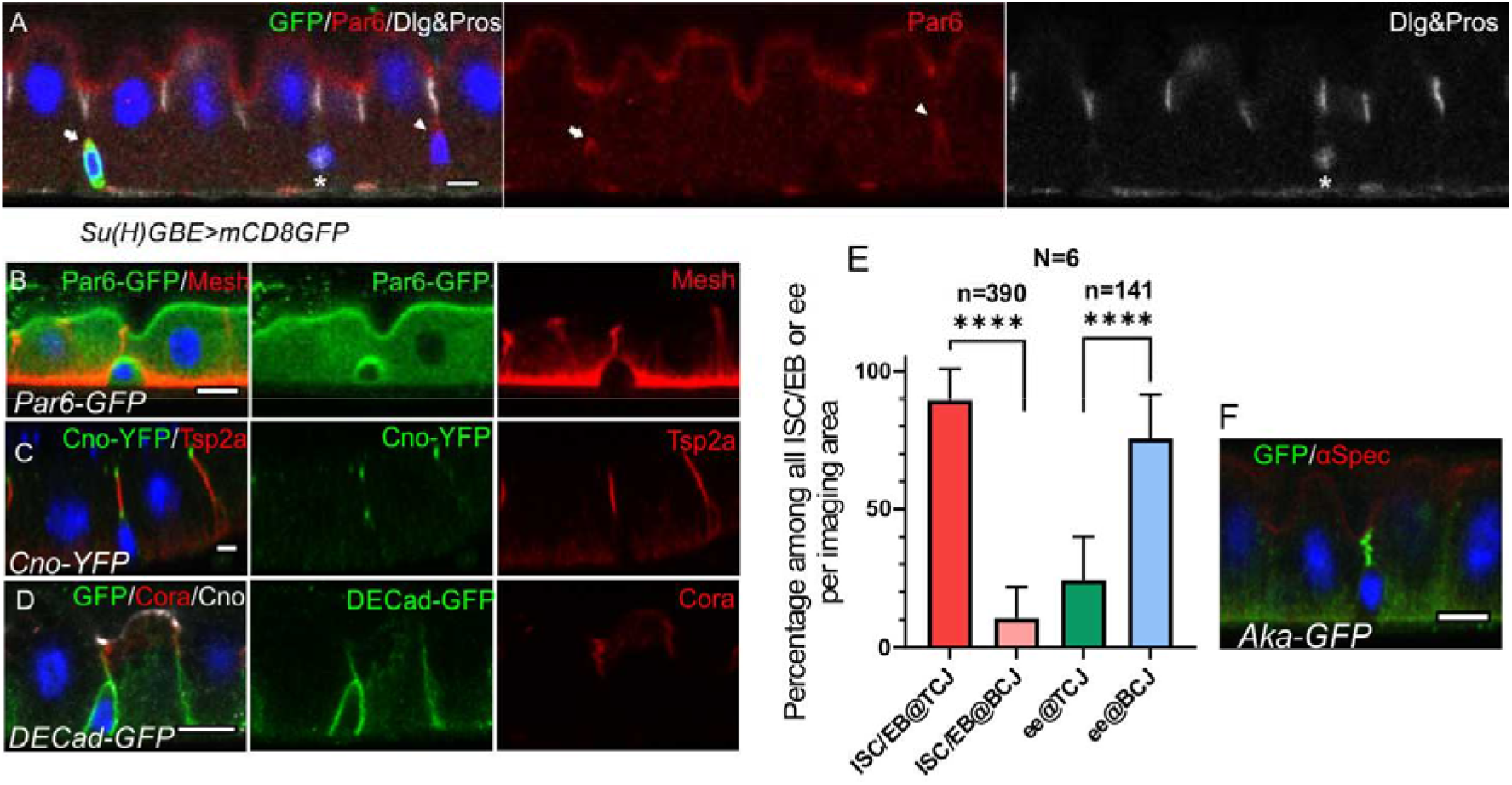
ISCs/early enteroblasts are polarised and reside underneath tri-cellular junctions. (A) Par-6 localises apically in ISCs and early enteroblasts. Su(H)GBE>mCD8-GFP expression (arrow) marks an early enteroblast, while the GFP-negative cell (arrowhead) is an ISC. Nuclear Prospero staining (white) marks an entero-endocrine cell (*) and cytoplasmic Dlg (greyscale) labels the septate junctions. (B) An ISC/early enteroblast expressing Par-6-GFP (green), which localises apically. Mesh (red) marks the septate junctions and basal labyrinth. (C) Canoe-YFP (green) localises to the marginal zone above the septate junctions (Tsp2a; red) in mature enterocytes. Although ISC/early enteroblasts do not form septate junctions, Canoe-YFP localises apically. (D) Adherens junctions (DE-cadherin-GFP; green) form throughout the cell-cell contacts between ISC/early enteroblasts and enterocytes. Coracle (red) marks the septate junctions; Canoe; white. (E) A graph showing the proportion of ISCs/early enteroblasts (ISC/EB) or entero-endocrine cells (ee) that localise beneath tri-cellular junctions (@TCJ) and bicellular junctions (@BCJ). Most ISCs/EBs localise beneath tricellular junctions where three enterocytes meet, whereas entero-endocrine cells mainly lie beneath bicellular junctions. DE-Cad-GFP expressing midguts were stained for GFP to mark cell contacts and Prospero to mark the ees. Cells with a low DNA content (~2n) were counted by imaging from the basal side. 390 ISC/early enteroblasts and 141 ee were scored in 6 guts. (F) An example of an ISC/EB beneath a tricellular junction marked by Anakonda-GFP (Aka-GFP; green). Scale bars, 5μm.

Canoe is a scaffolding protein that often links the actin cytoskeleton with junctional complexes(Boettner et al., 2003; Perez-Vale et al., 2021; Sawyer et al., 2009; Yu and Zallen, 2020). Canoe localizes to the most apical tip of the septate junctions in differentiated enterocytes, forming a thin belt separating the apical brush border (BB) from the lateral junctions (Figure 1 C and 1D). This location is equivalent to the marginal zone in other fly epithelia and the vertebrate marginal zone (VMZ) in mammalian epithelial cells(Tan et al., 2020; Tepass, 2012). Like Par-6, Canoe also localises apically in ISCs and early enteroblasts, which have no septate junctions (Figure 1B-1D, S1B and S1C). The apical localisation of Canoe and Par-6 in ISC/early enteroblasts indicates that they have some apical-basal polarity, although neither is required in the midgut.

The basal ISCs are found as single cells or as pairs of cells, which can either be two ISCs or an ISC and an enteroblast(de Navascues et al., 2012; Zhai et al., 2017). Imaging ECad-GFP flies from the basal side reveals that the ISC/early enteroblasts (Figure S1A) usually lie underneath the tri-cellular junctions (TCJ) between enterocytes rather than bi-cellular junctions (BCJ) (Figure 1E). In a side view of the epithelial layer, the ISCs/early enteroblasts sit immediately beneath the TCJs, marked by Anakonda-GFP (Figure 1F). This means that enteroblasts need to break through three mature enterocyte-enterocyte septate junctions to integrate into the epithelial layer and gain access to the gut lumen. More importantly, this integration process needs to be tightly regulated to ensure that gut barrier function is maintained.

### Integrating enteroblasts generate a pre-formed apical compartment (PAC)

Early enteroblasts express the Notch signalling reporter, Su(H)GBE-Gal4>mCD8GFP, but this is turned off as they differentiate(Micchelli and Perrimon, 2006; Ohlstein and Spradling, 2007; Rojas Villa *et al*., 2019). Enteroblasts that have just started to differentiate can therefore be identified as cells with larger nuclei, in which some GFP signal still perdures. The frequency of such early differentiating enteroblasts is very low under homeostatic conditions(Reiff and Antonello, 2019). We therefore exposed the flies to a 2 hour heat shock at 37 °C to induce minor damage and dissected them 1day later, which increases the number of dividing ISCs and differentiating Su(H)GBE>mCD8GFP positive enteroblasts with larger nuclear volumes (Figure S2A and S2B). This treatment causes a small increase in Tor kinase activity at the very anterior and posterior ends of the midgut as shown by phospho-4E-BP1 staining, but does not increase pERK staining, an indicator of EGFR/MAPK signaling (Figure S2C and S2D). Thus, the heat shock induces sufficient stress to trigger ISC divisions and enteroblast activation/differentiation but does not activate a regeneration process.

A proportion of the Su(H)GBE>GFP positive, differentiating enteroblasts and the older GFP-negative enteroblasts form a “bubble” like structure inside the epithelial layer (Figure 2A). This structure is surrounded by membranes, as shown by the localisation of the membrane associated protein, α-Spectrin (Figure 2A). The actin marker sqh∷Utrophin actin binding domain-GFP (sqh∷Utr-ABD-GFP), which labels the brush border of mature enterocytes, also localises to the lower portion of the “bubble” (Figure 1E). Furthermore, the actin-rich portion of the “bubble” is labelled by all other apical domain markers that we have examined, including Par-6, Myosin IA, Myosin7a, the actin cross-linking proteins, Fimbrin and Cheerio (Lcp1/Pls3 and Filamin in mammals), Rab11, a marker for the apical recycling endosome and Picot, a transmembrane protein with homology to anion co-transporter proteins (Figure 2B-2E, S2E and S2G). This strongly suggests that the lower portion of the “bubble” corresponds to a nascent apical domain. In support of this view, EM images reveal that this region forms a brush border (Figure 2F). Thus, the lower part of the “bubble” structure has all the features of a fully-developed apical domain, although it is not at the apical side of the epithelium.

**Figure 2.**
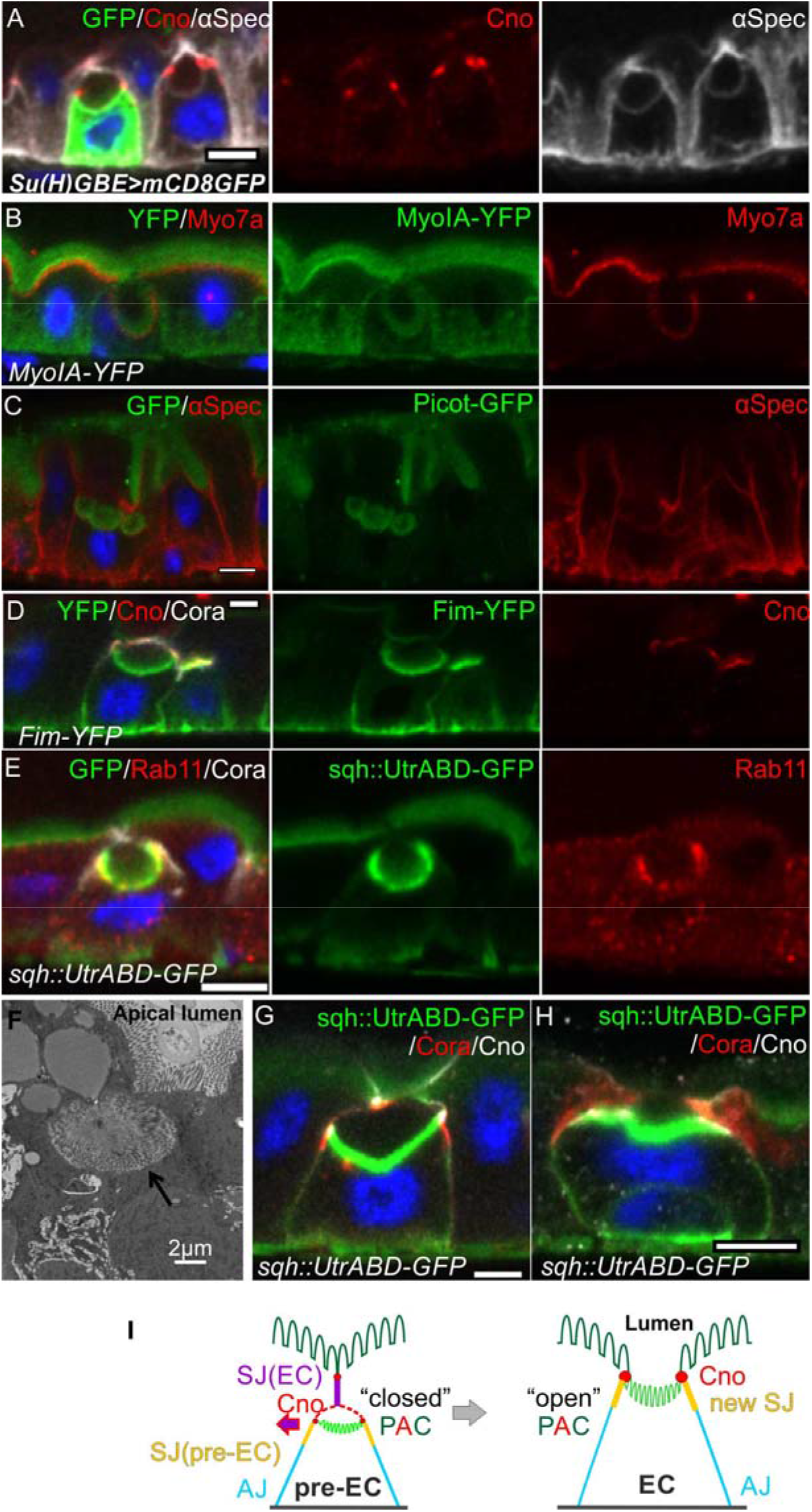
Integrating enteroblasts form an apical domain before reaching the gut lumen. (A) A transverse section of a *Su(H)GBE>mCD8GFP* midgut imaged 1 day after heat shock. Large spherical lumens surrounded by plasma membranes (α-spectrin; greyscale) have formed between the integrating enteroblasts and the overlying enterocytes. One enteroblast is still GFP-positive, indicating that it has recently been activated, whereas the other has lost Su(H)GBE>mCD8GFP (green) expression. Canoe (red) labels the apical corners of the enteroblasts. (B) The lumen-facing side of the enteroblast is marked by MyoIA-YFP (green) and Myo7a (red), which are markers for the enterocyte brush border and apical cortex respectively. (C) A GFP protein trap line in the transmembrane transporter Picot (green) labels the apical brush border in enterocytes and the lumen-facing membrane in an integrating enteroblast. Note that multiple lumens have formed between the enteroblast and the enterocytes above. Plasma membranes are labelled with α-spectrin (red). (D) Fimbrin-GFP (Fim-GFP; green) marks the apical cortex of an integrating enteroblast and the enterocyte brush border. Note that the enteroblast to the right has not yet formed a lumen but has Fimbrin, Canoe (red) and Coracle (greyscale) localised to its apical surface. (E) The actin binding domain of Utrophin (Sqh∷UtrABD-GFP; green) marks the enterocyte brush border and the apical side of an integrating enteroblast. The apical recycling endosome marker, Rab11 (red) also labels the apical region of the enteroblast. (F) A transmission electron micrograph showing that the lumen above an integrating enteroblast is surrounded by brush border microvilli. Scale bar, 2μm. (G) An integrating enteroblast with a closed pre-apical compartment (PAC) and lumen that lie below the septate junction between the overlying enterocytes. The cells express the actin marker, Sqh∷UtrABD-GFP, and are stained for Coracle (red) and Canoe (greyscale). (H) An integrating enteroblast stained as in (H) with an open lumen that is continuous with the gut lumen. (I) A model for enteroblast integration in which a “closed” PAC precedes an “open” PAC. Scale bars in A-E, G and H, 5μm.

To test whether the lumen facing the internal apical domain is continuous with the gut lumen, we examined the position of the brush border (sqh∷UtrABD-GFP), relative to the septate junctions (Cora) and the marginal zone (Canoe). In many cases, the internal apical domain lay beneath the septate junction connecting the overlying enterocytes, indicating that the lumen lies within the epithelium and does not connect to the gut lumen (Figure 2G, S2H, VideoS1). In other cases, the only septate junctions are the new junctions between the invaginating enteroblast and the junctions are the new junctions between the invaginating enteroblast and the neighbouring enterocytes and the lumen is continuous with the gut lumen (Figure 2H, S2I, VideoS2). Thus, the enteroblasts first form an apical domain and lumen within the epithelium. The septate junction between the neighbouring enterocytes then disappears, allowing the internal lumen to fuse with the gut lumen and the apical domain to reach the apical surface. Since these results indicate that the apical domain forms internally, we refer to this structure as a “Preformed Apical Compartment (PAC)” and refer to the cells with a PAC as pre-enterocytes (Figure 2I). In the “closed” PAC stage, a new set of septate junctions form beneath the septate junction between the neighbouring enterocytes. This means that the pre-enterocyte has septate junctions with its neighbours and a fully-developed apical brush border before it emerges on the apical surface, thereby maintaining the gut barrier during enteroblast integration (Figure 2I).

### Integrating enteroblasts initiate PAC formation by forming an apical AMIS

Imaging heat shocked flies expressing *sqh∷UtrABD-GFP* or *Fim-GFP* and stained for Canoe and Cora revealed three distinct stages of PAC formation. In the first stage, actin is diffusely distributed around the cell cortex, as it is in quiescent enteroblasts, but Canoe and Cora enriched apically (Figure 3A, S3A and S3C). Actin then becomes enriched apically in a slightly smaller domain than Canoe and Cora (Figure 3B, S3B and S3D). This co-localisation of actin and junctional proteins is reminiscent of the apical membrane initiation site (AMIS) in MDCK cells(Bryant *et al*., 2010).

**Figure 3.**
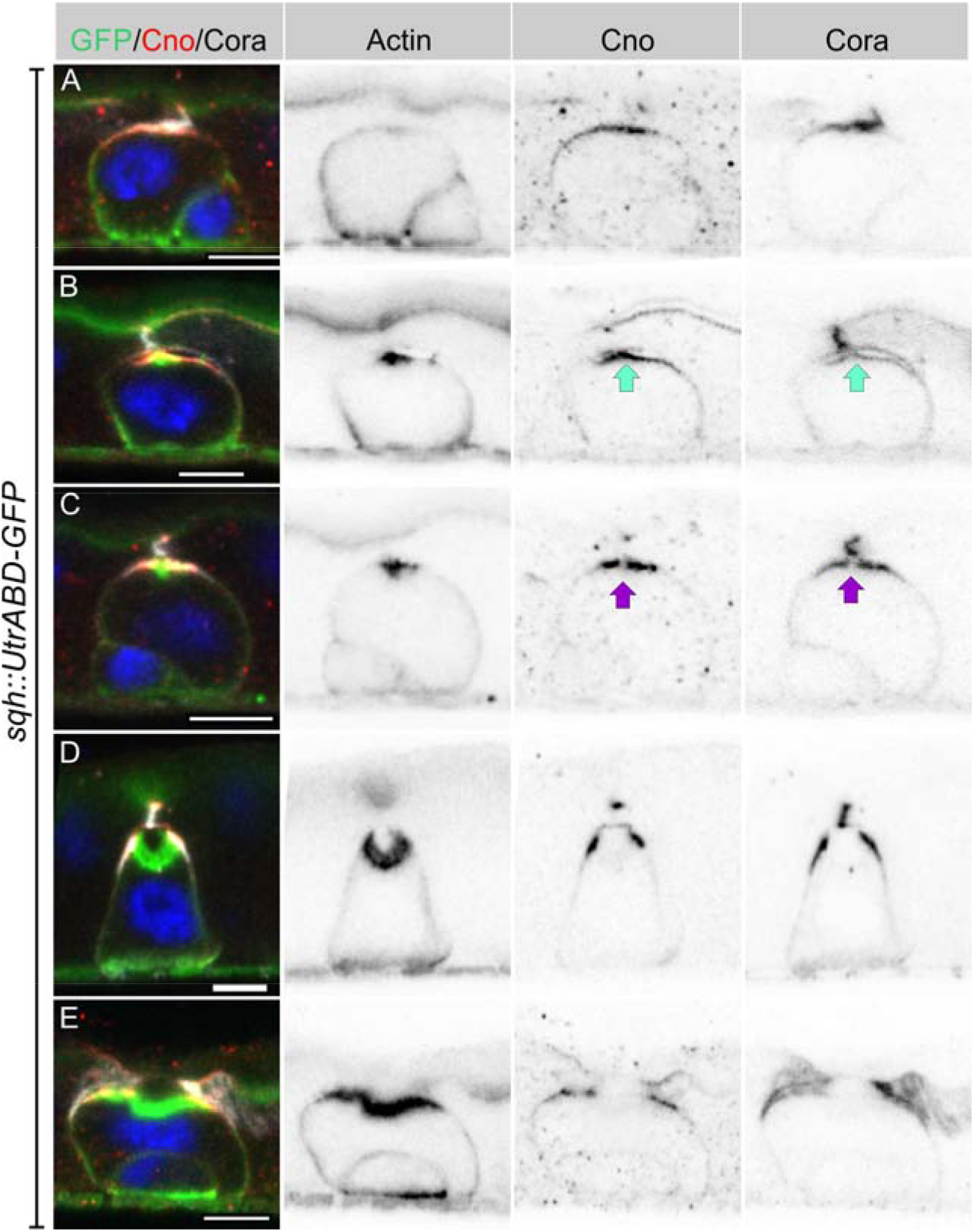
Integrating enteroblasts form an AMIS before forming a PAC. (A) When enteroblasts reach the level of the septate junction between the adjacent ECs, Canoe (red) and Cora (white) localise to the apical side of the enteroblast, whereas actin is still uniformly distributed around the enteroblast cortex (green, GFP staining in *sqh∷UtrABD-GFP* flies). (B) AMIS (green arrow) formation in an integrating enteroblast. Actin now localises to the apical side of the enteroblast in a smaller domain than Cora and Canoe. Note that the plasma membrane in the AMIS forming region has detached from the neighbouring ECs. (C) Actin concentrates in the centre of the AMIS, while Canoe and Cora are excluded from this region (purple arrow). (D) The actin enrichment enlarges to form a preformed apical compartment (PAC) below the septate junction between the neighbouring ECs. Cora localises to the septate junctions that are forming between the pre-enterocyte and mature enterocytes on either side. Canoe localises to the marginal zone above the septate junctions and is also enriched on the enterocyte membranes facing the PAC lumen. (E) The septate junction between the overlying enterocytes dissolves and allows the PAC to fuse with the gut lumen. Scale bars = 5μm.

Coincident with the apical enrichment of actin, a slight separation appears between the apical membrane of the enteroblast and the overlying enterocyte membranes, suggesting that fluid is being secreted into this space (Figure 3B). The actin staining then becomes concentrated in the centre of the apical domain, while Canoe and Cora are depleted from this region, creating a central actin-rich zone that lacks junctional proteins (Figure 3C). This small actin region then expands, bends inwards and becomes the PAC, whereas Cora localises to the newly-forming septate junctions on either side, with Canoe slightly more apical in the nascent marginal zone (Figure 3D). Finally, the pre-existing septate junction above the PAC disappears, and the PAC everts to form the apical domain (Figure 3E).

### AJs are cleared from the AMIS

The separation between the enteroblast and enterocyte membranes as the AMIS forms suggests that cell-cell adhesion must be modified at this stage. We therefore examined the localisation of Ecad-GFP and Armadillo during enteroblast integration. Early enteroblasts that have not reached the septate junction between the overlying enterocytes form adherens junctions (AJ) with the adjacent enterocytes all round their contacting surfaces (Figure 4A). At this stage, Cora is uniformly distributed around the cortex and Canoe is weakly enriched apically. Canoe and Cora become polarised apically when the integrating enteroblast contacts the enterocyte-enterocyte septate junction above, but the adherens junctions remain uniformly distributed (Figure 4B). As actin and Canoe become enriched apically to form the AMIS, adherens junctions are lost from this region and the enteroblast membrane separates from the overlying enterocytes (Figure 4C). Adherens junctions remain absent form this region as the PAC starts to form (Figure 4D, S4A and S4B). Thus, AMIS formation is associated with the loss of adherens junctions. This is presumably required to allow the enteroblast membrane to separate from that of the overlying enterocytes, as fluid is secreted from the AMIS to form the internal lumen.

**Figure 4.**
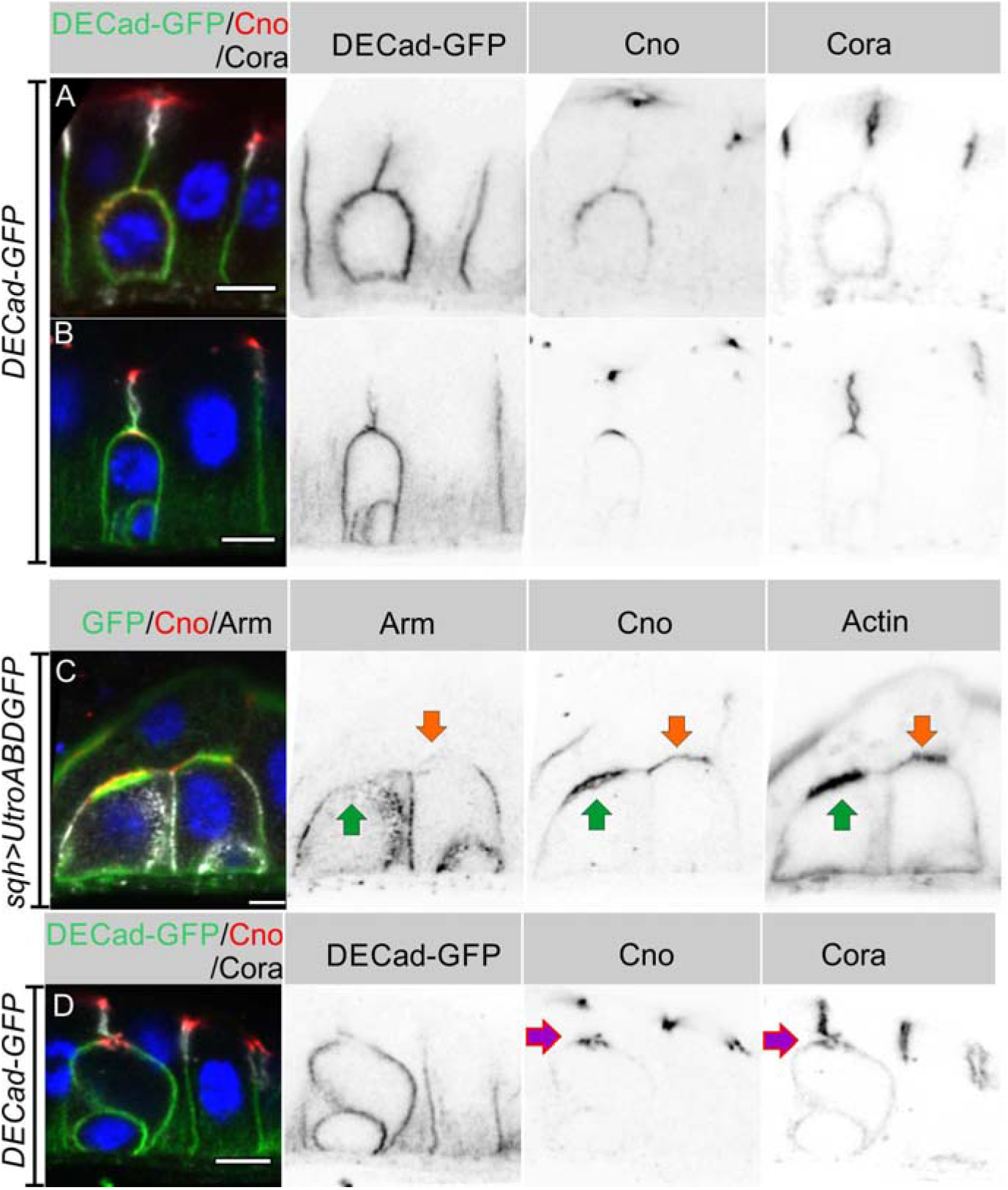
AJs are lost from the apical membrane during AMIS formation. (A) E-cadherin (green) localises all around the plasma membrane of integrating enteroblasts that have not yet reached the septate junction between the overlying enterocytes. Cora (white) is not polarised at this stage and Canoe (red) is only weakly enriched apically. (B) Canoe and Cora form apical caps in enteroblasts that have reached the septate junction between the overlying enterocytes, but E-cadherin is still uniformly distributed along the enterocyte-enteroblast cell contact sites. This enteroblast is at the stage when Canoe is polarised but actin is not (see Figure 3A, S3A and S3C). (C) AMIS formation (green arrow) in an integrating enteroblast. The Adherens junctions (Arm staining in white) disappear from the AMIS region. Canoe (red) localises to both the enteroblast and enterocyte membranes after their separation as seen in Figure 3B. The integrating enteroblast on the right (orange arrow) is at a slightly earlier stage before separation of the apical enteroblast membrane from the overlying enterocyte membranes. Actin (green) has started to accumulate apically, but at lower levels than in the left-hand enteroblast. (D) AJ (Ecad-GFP in green) is absent from the membrane around the pre-apical compartment (purple arrow) as it forms. Scale bars, 5μm.

### Sox21a is turned off at the enteroblast to pre-enterocyte transition

The transcription factor Sox21a is required for the differentiation of enteroblasts into enterocytes and can induce the precocious differentiation of quiescent enteroblasts when over-expressed, in part by activating the expression of another transcription factor, Pdm1(Chen *et al*., 2016; Meng and Biteau, 2015; Zhai *et al*., 2017). We therefore examined the expression of Sox21a during enteroblast integration and differentiation in heat-shocked flies expressing *sqh∷UtrABD-GFP*. The Sox21a antiserum labels the septate junctions, but this signal is non-specific, as it is still present in *sox21a* mutant cells (Figure S5A). Nuclear Sox21a is present at high levels in unpolarised enteroblasts, which probably correspond to the oval-shaped enteroblasts described by Chen. et.al (2016), but the levels are significantly lower in polarised enteroblasts (Figure 5A, 5B and 5D). Sox21a is no longer detectable above background in the nuclei of pre-enterocytes that have formed a PAC, with a nuclear intensity similar to that in the neighbouring Sox21a-negative enterocytes (Figure 5B and 5D). Indeed, the pre-enterocytes are similar to enterocytes at the transcriptional level, as they also express the enterocyte marker, Pdm1 (Figure 5C). Sox21a levels are therefore inversely correlated with the enteroblast differentiation and integration: unpolarised enteroblasts have high Sox21a, polarised enteroblasts with an AMIS have lower levels, and pre-enterocytes with a PAC are Pdm1+ and Sox21a-like the neighbouring enterocytes (Figure 5E). However, down-regulation of Sox21a is not essential for enteroblast to pre-enterocyte differentiation, since enteroblasts expressing *UAS-Sox21a* under the control of *esg-Gal4* still from a PAC (Figure S5B).

**Figure 5:**
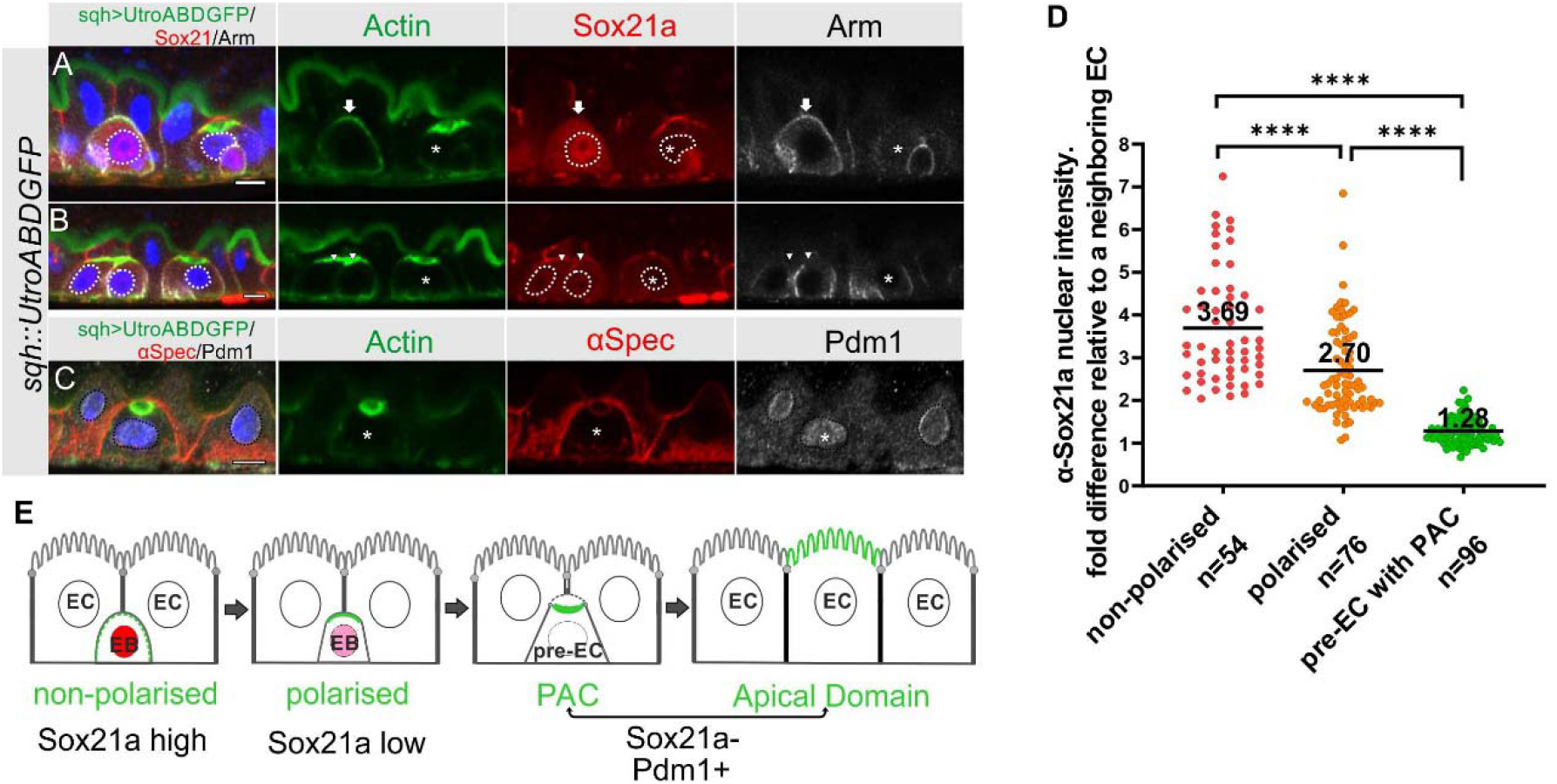
Sox21a levels fall as the PAC forms during integration. (A-B) Sox21a (red) is present at high levels in the nuclei of enteroblasts in which actin (Utr-ABD-GFP; green) is not yet polarised apically (arrow in A), is lower in the nuclei of enteroblasts with polarised actin (arrowheads in B). Pre-enterocytes that have formed a PAC (asterisks in A and B) lack nuclear Sox21a. The nuclei are outlined by white dashed lines. Note that the Adherens junctions (Armadillo; white) still extend around the apical membrane of the enteroblast with unpolarised actin, but this signal has disappeared in the enteroblasts with apical actin (AMIS stage, arrowheads in B) and the pre-enterocytes with a PAC (*). The anti-Sox21a antiserum labels the septate junctions, but this is non-specific staining as it is still present in *Sox21a* mutant flies (see Figure S5A). (C) Pdm1+ (white) is expressed in a Pre-enterocyte with a PAC (asterisk); actin in green and αSpec in red. Scale bar=5μm. (D) Graph showing the levels of nuclear Sox21a staining relative to neighbouring enterocytes at different stages of enteroblast integration. The horizontal lines indicate the median values, which are significantly different by a 2-tailed t test among three groups (**** p< 0.0001). n, the number of EB/neighbouring EC pairs. (E) Diagram showing the levels of nuclear Sox21a and Pdm1 during enteroblast integration.

Combining all our analyses, we propose the following steps in enteroblast differentiation (Figure 6). When an enteroblast is activated by high Sox21a expression, it starts to grow in size but becomes unpolarised, with adherens junctions all over its contacting surface and uniform cortical actin. Once the enteroblast reaches the septate junction between the neighbouring enterocytes, Canoe and Cora become localised apically. Shortly afterwards, the apical membrane initiation site forms, with the apical localisation of F-actin and the removal of DE-cadherin from this region. This coincides with the down-regulation of Sox21a and the opening of a space between the plasma membranes of the enteroblast and the overlying enterocytes, suggesting that the apical membrane is secreting fluid between the cells. As the cell differentiates into a pre-enterocyte, F-actin segregates from Canoe and Coracle into the centre of the AMIS to form the nascent PAC. The PAC then expands and new septate junctions (marked by Coracle, Mesh (Figure S2G) and Tsp2A (Figure S4A) form between the enteroblast and the adjacent enterocytes, sealing the fluid-filled lumen above the PAC. This coincides with the localisation of Canoe to the marginal zone above the new septate junctions and in the enterocyte membrane covering the PAC. By this stage, the pre-enterocyte no longer expresses Sox21a and has activated Pdm1. Finally, the internal PAC fuses with the gut lumen and forms an apical domain, with new septate junctions replacing the old junction that covered the PAC (Figure 6).

**Figure 6.**
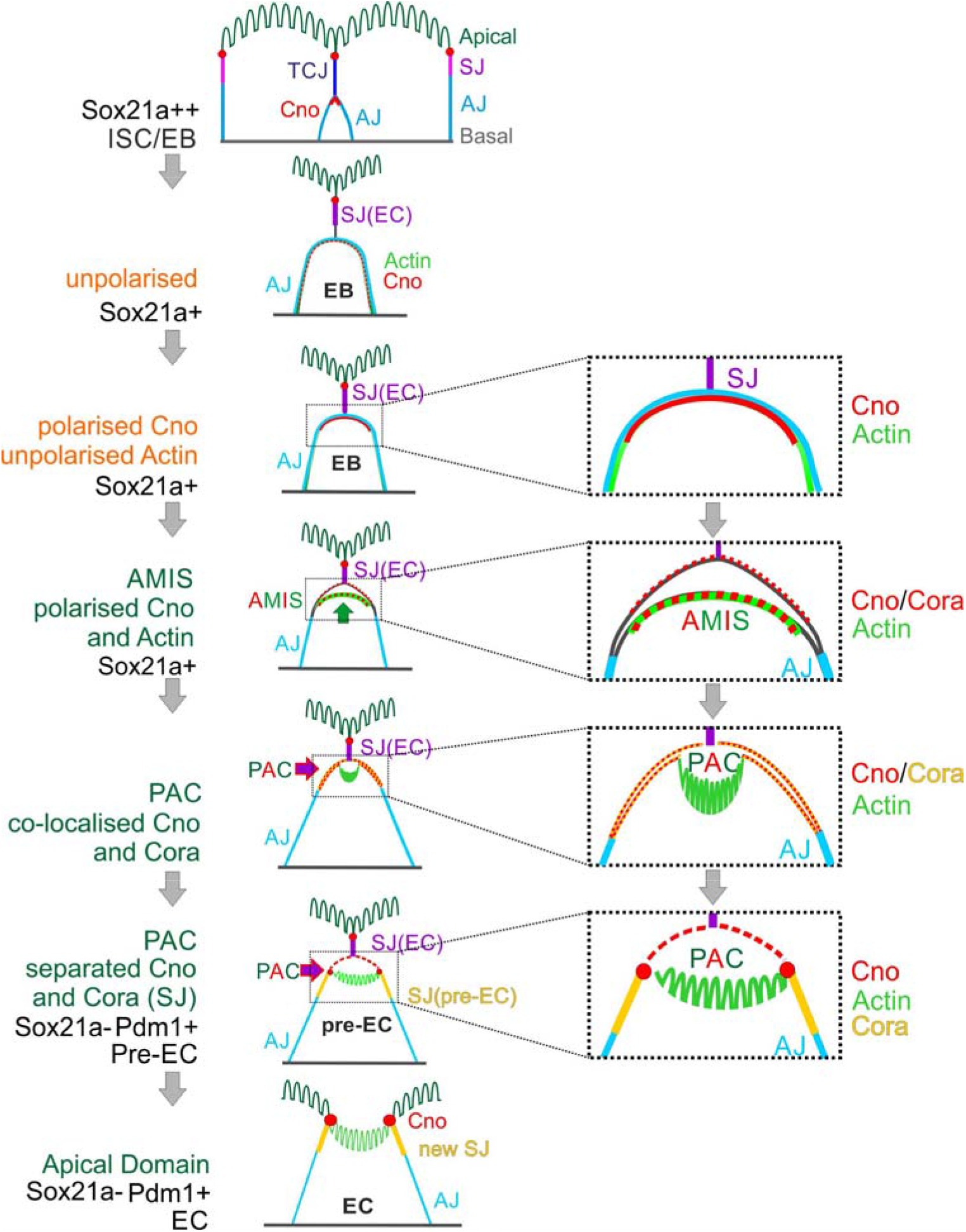
Diagram of the steps in enteroblast integration.

### Canoe is not required for PAC formation or cell polarity

Canoe provides a useful marker to follow enteroblast integration, as it is the first protein detected apically as the enteroblast polarises and it then labels the AMIS and the outer rim of the PAC. During AMIS and PAC formation, we noticed that Canoe also localises to the enterocyte membranes that separate from the apical membrane of the enteroblast and cover the internal lumen (orange arrow in Figure 4C, yellow arrow in Figure 7A). Since Canoe normally localises to the marginal zone above the septate junctions in enterocytes, this suggests that enterocytes respond to the presence of an invaginating enteroblast/pre-enterocyte by re-localising Canoe. To confirm that this is the case and to test whether Canoe plays a functional role in enteroblast integration, we generated positively marked clones homozygous for the null allele *Canoe*^R10^ using the MARCM system(Lee and Luo, 2001; Sawyer *et al*., 2009). When one of the adjacent enterocytes is mutant for *Canoe*, Canoe is lost from the enterocyte membrane covering that side of the internal lumen, confirming that this signal normally comes from the enterocyte (Figure 7B). Loss of Canoe from the pre-enterocyte has no obvious effect on Canoe localisation, presumably because both the pre-enterocyte and the neighbouring enterocytes contribute to its localisation at the marginal zones above the new septate junctions (Figure 7C, S6A and S6B). In both cases, however, the PAC forms normally, suggesting that Canoe is not required for enteroblast integration or polarisation. To rule out the possibility that the lack of a phenotype was due to residual function of the *Canoe*^R10^ mutant, we used CRISPR to generate a 14bp deletion and premature stop codon in DIL domain of Canoe. MARCM clones of this allele, *Canoe*^jc^, lack Canoe staining, but are otherwise wild-type, confirming that Canoe is not required for enteroblast integration (Figure 7D, 7E, S6C and S6D).

**Figure 7.**
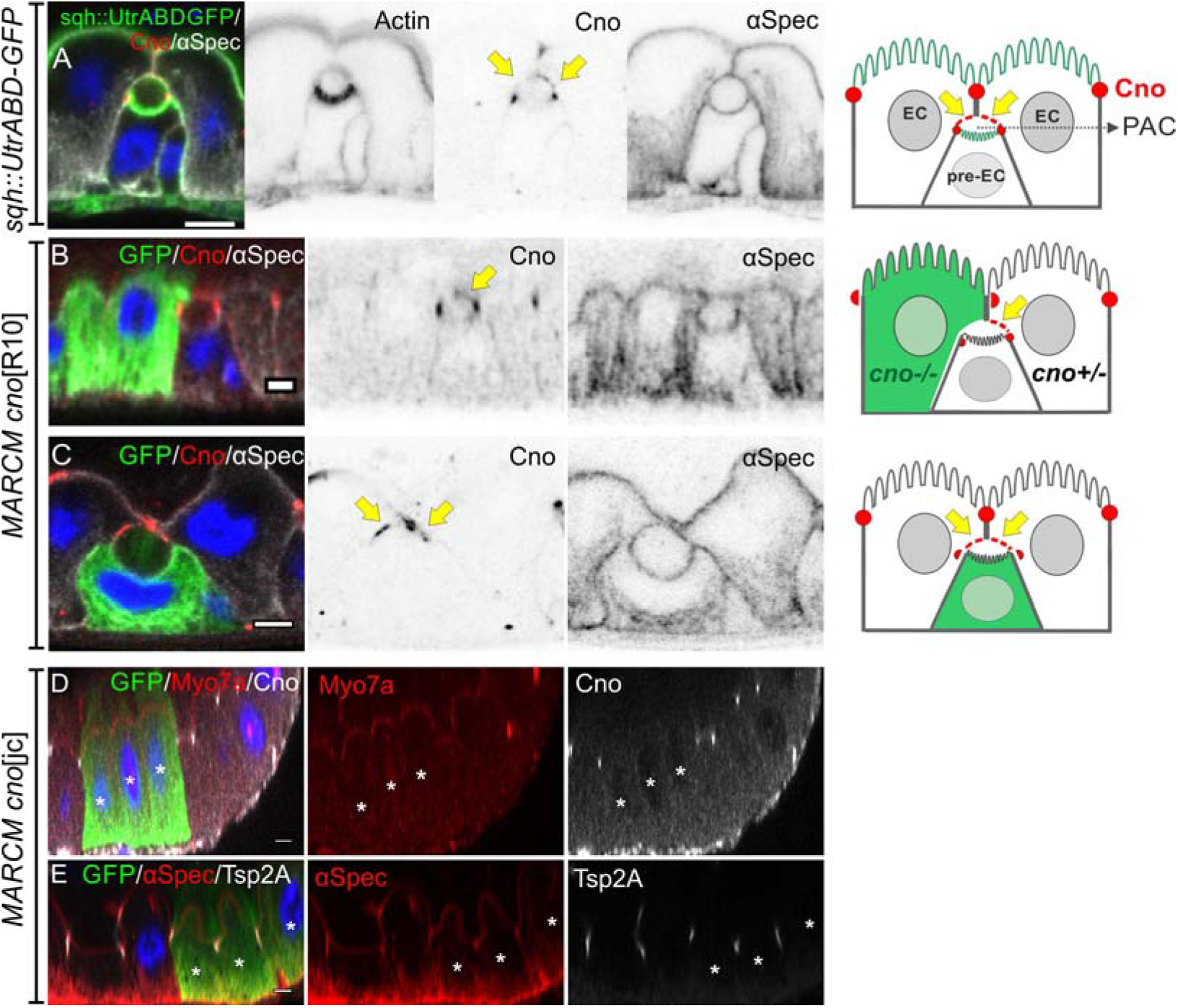
Canoe is not required for PAC formation or enterocyte polarity. (A) Canoe (red) localises to the enterocyte membranes (yellow arrows) that face the lumen above an integrating pre-enterocyte, and to the marginal zone above the newly formed septate junctions between the pre-enterocyte and neighbouring enterocytes. Sqh>UtrABD-GFP labelling of Actin in green and α-spectrin in greyscale. (B) An enteroblast integrating between a *canoe*^R10^ mutant enterocyte (green) and a heterozygous enterocyte. Canoe (red) is lost from the roof of the lumen on the side with the mutant enterocyte, but still marks the roof on the side with a non-mutant enterocyte (yellow arrow). (C) A *canoe*^R10^ mutant pre-enterocyte (green) integrating between two heterozygous enterocytes. The PAC still forms normally in the absence of Canoe (red). α-Spectrin in greyscale. (D-E) MARCM clones of *canoe*^JC^ homozygous mutant cells stained for Myo7a (red; D), ◻-spectrin (red; E) and Tsp2A (greyscale; E). The mutant cells form normal apical domains and septate junctions in the absence of Canoe. * marks the homozygous mutant cells. Scale bar, 5μm.

### Septate junctions are required for normal PAC formation

The smooth septate junctions of the *Drosophila* midgut are formed by several transmembrane proteins, including Mesh, Tsp2A and Snakeskin (Ssk), which are specifically expressed in endodermal tissues(Furuse and Izumi, 2017). The apical localisation of these proteins coincides with the formation of the AMIS, while PAC formation correlates with their lateral displacement, raising the question of whether they play role in these processes.

We examined the effects of loss of *mesh* by generating positively-marked MARCM clones of *mesh*^f04955^ and a new *mesh* null allele, *mesh*^R2^, which we identified as a second hit on the *canoe*^R2^ chromosome that abolishes Mesh antibody staining (Figure 8A). *mesh*^R2^ and *mesh*^f04955^ homozygous cells never integrate fully into the epithelial layer and remain below the septate junctions of the overlying enterocytes, presumably because they cannot make septate junctions of their own (Figure 8A-8C). Some of the mutant cells contain one or more circular, actin-rich structures that are labelled by α-spectrin and the apical domain proteins, Par-6, Myo7a and Rab11 (Figure 8C, 8E, 8G and 8H). These structures resemble PACs, but do not face the external space between the enteroblast and enterocytes, instead forming inside the cells. Cells with an internal PAC account for 12% of the *mesh*^R2^ enteroblasts. 81% of differentiating *mesh* mutant enteroblasts, based on their size and ploidy (>8n), do not appear to be polarised, with actin and β_H_-Spectrin diffusely distributed around the contacting cell surface (Figure 8D and 8J). The remaining mutant cells (7%) are polarised, with actin, Par-6 and Rab11 localised apically (Figure 8E, 8F and 8J).

**Figure 8.**
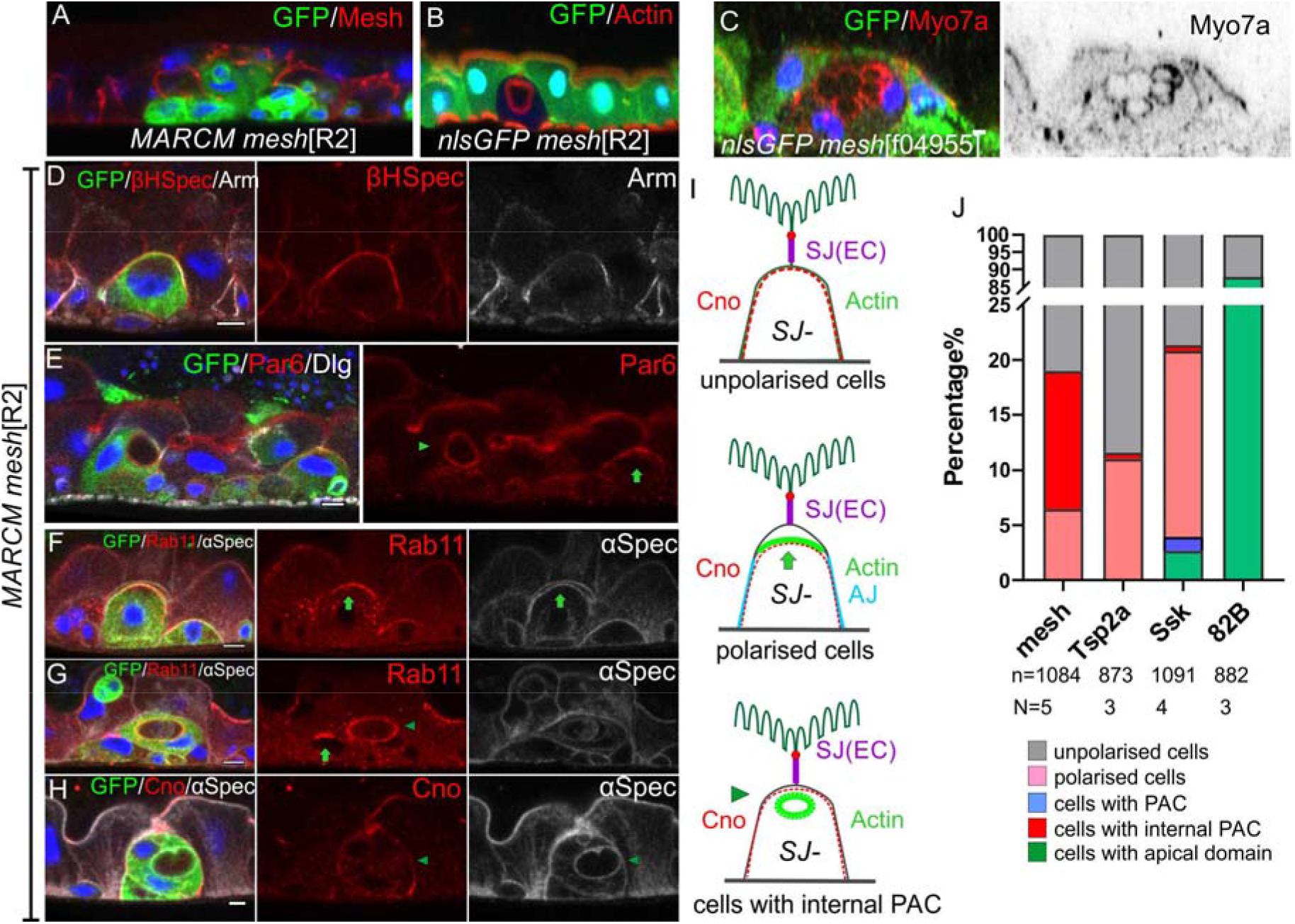
*mesh* mutants fail to form a PAC or form an internal PAC. (A) A *mesh*^R2^ homozygous MARCM clone marked by GFP (green). The mutant cells lack Mesh staining (red), fail to make septate junctions and do not reach the gut lumen. (B-C) *mesh*^R2^ (B) and *mesh*^f04955^ (C) homozygous mutant cells marked by the loss of GFP (green). The mutant cells form internal PACs that are marked by actin (red in B) and Myo7a (red in C). The cell in C has formed multiple internal PACs. (D) β_H_-spectrin (red) localises around the plasma membrane of *mesh*^R2^ mutant enteroblasts (green), indicating that they do not polarise normally, but adherens junctions (Arm, greyscale) are still down-regulated at the apical surface. (E) *mesh*^R2^ mutant enteroblasts (green) stained for Par-6 (red). Par-6 localises to the apical membrane (green arrow) in the enteroblast on the right, but localises around the internal PAC in the enteroblast on the left (green arrowhead). (F) A *mesh*^R2^ mutant enteroblast stained for a-spectrin (greyscale) and Rab11 (red), which localises apically (green arrow). The a-spectrin staining reveals that a space has formed between the apical side of the integrating enteroblast and the neighbouring enterocytes. (G) A *mesh*^R2^ mutant enteroblast stained for α-spectrin (greyscale) and Rab11 (red) that has formed an internal PAC. Rab11 decorates the surface of the internal PAC (green arrowhead). Note the younger, mutant enteroblast on the left (green arrow) localises Rab11 apically. (H) A *mesh*^R2^ mutant enteroblast (green) with an internal PAC stained for Canoe (red) and a-spectrin (greyscale). Canoe does not localise to the internal PAC, which has no junctions. Scale bars = 5μm. (I) Diagrams showing the distributions of Canoe, actin and adherens junctions in the three phenotypic classes of *mesh*^R2^ mutant enteroblasts. (J) Percentages of *mesh*, *Tsp2a* and *ssk* mutant enteroblasts in each of the classes in (I) compared to a wild-type control (FRT82B clones) based on actin in mutant cells (representative images in Figure S7E and S7F).

Mutants in other septate junction proteins have similar phenotypes. Only 0.6% *Tsp2a* mutant cells have an internal PAC and 11% still polarise, with the rest showing only very weak signs of polarity (Figure 8J, S7E, S7F, S7K and S7L). Similarly, 78.6% of *ssk* mutant cells do not polarise, 17% polarise and 0.5% form an internal PAC (Figure 8J, S7H and S7I). However, a small fraction of *ssk* mutant cells form a normal PAC (1.3%) or even integrate into the epithelial layer (2.6%) (Figure 8J, S7G and S7J). By contrast, more than 80% of the cells in control *FRT82B* MARCM clones develop a mature apical domain, while the remaining ~10% lack apical actin and are presumably ISCs or early enteroblasts (Figure 8J). Mutants in each septate junction protein disrupt the localisation of the other septate junction components, as previously reported for the embryonic and larval gut (Figure S7A-S7C)(Izumi *et al*., 2021; Izumi *et al*., 2016; Izumi *et al*., 2012). The differences between the spectrum of phenotypes observed in *mesh*, *Tsp2a* and *ssk* mutants cannot therefore be explained by their distinct effects on septate junction formation and may reflect other functions of these proteins, such as the regulation of the Yki pathway(Chen et al., 2020; Izumi et al., 2019; Xu et al., 2019).

About 10% of cells mutant for septate junction proteins still polarise actin, Par-6 and Rab11 apically (Arrows in Figure 8E-8G, 8H, Fig S8A-S8E). Adherens junctions are also removed from this region and the apical membrane detaches from the overlying wild-type enterocytes (Figure 8D and 8F). All these features suggest that some enteroblasts can still polarise to form an AMIS-like structure in the absence of septate junctions. These results indicate that septate junctions are not absolutely required for apical-basal polarity in integrating enteroblasts, although they are essential for normal PAC formation. Surprisingly, Canoe is not enriched apically in this class of enteroblasts, although its apical localisation is unaffected in *mesh* mutant ISCs or early stage enteroblasts (Figure 8H, S8A-S8E). Thus, the septate junction proteins are not required for the initial polarisation of ISCs and quiescent enteroblasts, but they are required for the apical-basal polarisation that gives rise to the AMIS in most, but not all cells. This contrasts with the role of integrin adhesion complex, which is required for apical-basal polarity in ISCs, quiescent enteroblasts and activated enteroblasts (Figure S8F-S8H).

## Discussion

Enteroblasts perform an essential role in gut homeostasis by providing a pool of quiescent precursor cells that can rapidly integrate into the epithelium in response to damage or growth signals to replace or increase the number of enterocytes. In order to do this, the enteroblasts need to insert themselves between existing enterocytes, while maintaining the barrier function provided by the smooth septate junctions. Our results reveal that they do this by an unexpected mechanism, in which they form an apical domain inside the epithelial layer, immediately beneath the septate junction between the overlying enterocytes. At the same time, the enteroblast/pre-enterocytes form new septate junctions with their neighbours. This therefore provides an elegant solution to the problem of integrating into the epithelium without disrupting the barrier, by forming junctions between the enteroblast/pre-enterocyte and its neighbours before the septate junctions above the integrating cell are removed. This also means that the pre-enterocyte already has a fully differentiated apical domain with a brush border when it reaches the gut lumen, so that it can immediately assume its function in nutrient absorption.

Quiescent enteroblasts are induced to differentiate in response to damage or growth signals from the visceral muscle, trachea, enterocytes and ISCs(Jiang and Edgar, 2011; Miguel-Aliaga *et al*., 2018; Nászai *et al*., 2015). This leads to the down-regulation of *esg* and the Dronc caspase, which maintain enteroblasts in the undifferentiated state, and the up-regulation of factors that drive differentiation, such as Zfh2, Sox100b, Sox21a and Pdm1(Amcheslavsky et al., 2020; Jasper, 2020; Jiang et al., 2016; Meng and Biteau, 2015; Meng et al., 2020; Rojas Villa *et al*., 2019). These changes induce the enteroblasts to increase in size under the control of the Insulin, TOR and EGFR signalling pathways and to endo-replicate and become polyploid(Xiang *et al*., 2017). Activated enteroblasts has also been found to develop basal, actin-rich protrusions and to migrate occasionally to other regions of the epithelium (Antonello *et al*., 2015; Rojas Villa *et al*., 2019). Unlike these earlier studies, which imaged the midgut from the basal side, we have analysed the apical-basal axis of the enteroblasts as they are activated. This reveals that growing enteroblasts go through a phase where they are unpolarised, in contrast to ISCs and quiescent enteroblasts, which are polarised and localise Canoe and Par-6 apically. These unpolarised enteroblasts are likely to correspond to the protrusive, migratory state of activated enteroblasts reported by Antonella et al. (2015), indicating that growing enteroblasts go through a mesenchymal stage. Enteroblasts then re-polarise once they reach the septate junctions between the overlying enterocytes. Thus, the adult enteroblasts go through a similar set of transitions to the embryonic midgut precursors, which undergo an epithelial to mesenchymal transition when they delaminate from the primary epithelium and become migratory, before re-polarising in contact with the visceral mesoderm to form the embryonic midgut epithelium(Campbell et al., 2011; Pitsidianaki *et al*., 2021; Tepass and Hartenstein, 1994).

The re-polarisation of growing enteroblasts upon reaching the enterocyte-enterocyte septate junction occurs in two steps, with Canoe and septate junction proteins appearing at the apical membrane first, followed by F-actin. The apical localisation of actin coincides with the clearance of adherens junctions from the apical membrane and is followed by the appearance of a space between the apical membrane of the enteroblast and the overlying enterocytes. This suggests that the enteroblast is secreting fluid into the extracellular space between it and the enterocytes to separate the cell membranes, which are no longer held together via the E-cadherin adhesion. This may be driven by apical exocytosis of vesicles or the activity of water and ion channels. Actin and apical membrane markers such as Par-6, Picot and Rab11 then start to concentrate in the centre of the AMIS, while septate junction proteins are excluded from this region and start to form new junctions with the neighbouring enterocytes laterally, thereby creating a seal on either side of the forming lumen. The central actin-rich region then expands to form the pre-apical compartment and often invaginates inwards, perhaps due to the fluid pressure of the lumen above.

The steps in the formation of the pre-apical compartment bear several similarities with cyst formation in mammalian epithelial cultures in 3D, which also involves the development of an internal lumen at sites of cell-cell contact. In MDCK cysts, for example, the first sign of lumen development is the formation of an apical membrane initiation site, which is marked by the co-localisation of tight junction proteins, such as Cingulin and Occludin with the apical marker, Podocalyxin (Bryant *et al*., 2010). The AMIS then develops into a pre-apical patch as Cingulin and Occludin segregate away from the apical factors to form lateral tight junctions that seal the lumen, mirroring the behaviour of septate junction proteins in *Drosophila* enteroblasts(Blasky *et al*., 2015; Mangan *et al*., 2016). Similarly, the formation of an internal lumen in MDCK cysts is preceded by the loss of Cadherin from the contacting cell-cell surfaces, as it is in *Drosophila* enterocytes (Ferrari et al., 2008). Furthermore, junctional proteins are required to define the site of apical secretion in both systems, as knockdown of Cingulin blocks the formation of a single lumen in MDCK cysts and loss of Mesh or Tsp2a prevents the development of an external lumen in the *Drosophila* midgut (Mangan *et al*., 2016).

Despite these similarities, the spatial cues that determine where the AMIS forms appear to be different. In mammalian epithelial cysts, the site of the AMIS is defined by the position of the midbody formed during the last cell division, whereas enteroblasts are postmitotic and are derived from intestinal stem cell divisions that may have occurred several days earlier(Mangan *et al*., 2016; Rojas Villa *et al*., 2019). The most likely cue for the repolarisation that creates the AMIS in *Drosophila* enteroblasts is the septate junction between the overlying enterocytes, as enteroblasts do not polarise until they reach this junction. 80-90% of enteroblasts mutant for septate junction components fail to polarise at this stage, suggesting that the septate junction proteins may play a role in sensing when the integrating enteroblast contacts the enterocyte-enterocyte septate junction. However, this cannot be by forming three-way septate junctions with the existing septate junctions, because the enteroblast only forms septate junction later and in a more lateral position.

Another important difference between the two systems is that mammalian epithelial cysts are symmetric, with the cells on both sides of the lumen forming apical domains, whereas only the integrating enteroblast forms an apical domain in the fly midgut. The overlying enterocytes do not form a normal lateral domain over the lumen, however, as β_H_-Spectrin and Canoe localise to the enterocyte cortex in this region, which differs from their normal localisation in enterocytes to the apical domain and to the marginal zone above the septate junctions, respectively. This highlights the fact that the enterocytes above an integrating enteroblast are not passive bystanders but respond to the presence of the enteroblast and perhaps even also facilitate its integration. For example, the enterocytes must disassemble the septate junction above a pre-enterocyte for it to reach the gut lumen. This suggests that the enterocytes receive cues from the invaginating enteroblast that produce specific responses at each stage of the process, although the nature of these signals is not known.

The development of an enterocyte from an ISC involves first the loss of the polarity shown by ISCs and quiescent enteroblasts and then the re-establishment of polarity as the AMIS forms. Mutants in components of the integrin adhesion complex, such as Talin and Kindlin, disrupt both the initial polarity seen in ISCs and quiescent enteroblasts and the repolarisation during AMIS formation. However, the formation of the AMIS and PAC also requires septate junction proteins. Thus, differentiating enteroblasts only require a basal cue to establish their initial apical-basal polarity, whereas the formation of the pre-apical compartment also requires a junctional cue. The septate junctions are not necessary for apical domain formation per se, however, as *mesh* mutant enteroblasts form a full-developed apical domain with a brush border inside the cell. This suggests that septate junctions define the site of apical domain formation by delimiting the region where apical membrane proteins are secreted to assemble the brush border, but do not control the process of apical domain formation directly.

The internal apical domains in *mesh* mutant cells resemble the apicosomes observed in single human pluripotent stem cells in culture, which then fuse with the plasma membrane to form an extracellular lumen after cell division(Taniguchi et al., 2017). The apicosome is thought to form by the intracellular fusion of exocytic vesicles, and a similar mechanism may give rise to the internal apical domains in *mesh* mutant cells. If this is the case, it suggests that the apical domain can self-assemble and the polarity system merely functions to position where this domain forms. We cannot rule out the alternative possibility, however, that the internal apical domains in *mesh* mutants form by endocytosis of the AMIS region, as has been proposed for the intracellular vesicles with brush borders observed in multi-villus inclusion disease(Engevik et al., 2019; Engevik et al., 2021).

The polarisation of the midgut epithelium does not require any of the canonical epithelial polarity factors that polarise all *Drosophila* epithelia derived from the ectoderm and mesoderm, and instead depends on basal cues from adhesion to the extracellular matrix(Chen and Sayadian, 2018). Our observations suggest that a possible reason for this difference is that integrating enteroblasts polarise in a basal to apical direction and form an apical membrane without having a free apical surface. This means that they cannot use the apical polarity cues that trigger the polarisation of all other *Drosophila* epithelia. Thus, the unusual polarity system in the midgut may be a consequence of the way the tissue is built and maintained by basal stem cells. It will be interesting to determine if this is also the case for other epithelia with similar cellular arrangements.

## Materials and methods

### *Drosophila melanogaster* stocks

#### Fluorescently tagged protein lines

sqh∷UtrABD-GFP (gift from Thomas Lecuit, The Developmental Biology Institute of Marseille (IBDM), France), Par-6-GFP (Wirtz-Peitz et al., 2008), MyoIA-YFP (Lowe et al., 2014)(Kyoto DGRC #115611), Canoe-YFP (Lowe *et al*., 2014)(Kyoto DGRC#115111), Fimbrin-GFP (Bloomington #51562), Cheerio-YFP (Kyoto DGRC#115514), Picot-GFP (Bloomington #50822), DECad-GFP (gift from Yang Hong, University of Pittsburgh, USA), Anakonda-GFP(Byri et al., 2015) (gift from Stefan Luschnig, University of Münster, Germany).

#### Mutant stocks

FRT19A *Tsp2A*^1-2^, *Tsp2A*^3-3^, *Tsp2A*^2-9^ (Izumi *et al*., 2016)(gifts from Mikio Furuse, National Institute for Physiological Sciences, Okazaki, Japan), FRT82B *mesh*^R2^ (this paper), FRT82B *mesh*^f04955^ (Kyoto DGRC #114660), FRT80B *ssk*^1^ and *ssk*^2^ (Chen *et al*., 2020)(gifts from Tony Ip, University of Massachusetts Medical School, Worchester, MA, USA), FRT82B *canoe*^R10^ (Sawyer *et al*., 2009)(gift from Mark Peifer, University of North Carolina, Chapel Hill, NC, USA), FRT82B *canoe*^JC1^ (this study), FRT2A *Sox21a*^6^ (Zhai et al., 2015)(Bloomington #68389), FRT2A *Fit1*^18^ *Fit2*^83^, *rhea*^79a^ (Klapholz et al., 2015)(gifts from B. Klapholz and N. Brown, Department of Physiology, Development and Neuroscience, University of Cambridge, UK), UAS responder lines: UAS-Sox21a (Zhai *et al*., 2015)(Bloomington #35748), Su(H)GBE>GFP: Su(H)GBE-Gal4, UAS-mCD8GFP esg^ts^: esg-Gal4, UAS-mCD8GFP, tub>Gal80^ts^

Delta^ts^: Delta-Gal4, UAS-mCD8GFP, tub>Gal80^ts^

The following stocks were used to generate (positively-labelled) MARCM clones (Lee and Luo, 2001):

MARCM FRT82B: y w, UAS-mCD8∷GFP, Act5C-GAL4, hsFLP[1];; FRT82B tubP-GAL80.

MARCM FRT19A: w, hsFLP, tubP-GAL80, FRT19A;; tubP-GAL4, UAS-mCD8∷GFP/TM3, Sb.

MARCM FRT2A: hsFLP[1]; tubP-GAL4, UAS-mCD8∷GFP/CyO, GFP; FRT2A tubP-GAL80 (gift from B. Klapholz and N. Brown).

MARCM FRT80B: hsFlp[1]; tubP-GAL4, UAS-mCD8∷GFP/CyO, GFP; FRT80B tubP-GAL80 (generated by Dr. Mihoko Tame)

Negatively marked *mesh* mutant clones were generated using the following stock: esg-GAL4, UAS-FLP, tubP-GAL80^ts^/CyO; FRT82B nlsGFP (gift from Dr. G. Kolahgar, Department of Physiology, Development and Neuroscience, University of Cambridge, UK).

### Stock maintenance

Standard procedures were used for *Drosophila* husbandry and experiments. Flies were reared on standard fly food supplemented with live yeast at 25 °C. For the conditional expression of UAS constructs (e.g., overexpression of Sox21a) in adult flies, parental flies were crossed at 18 °C and the resulting offspring reared at the same temperature until eclosion. Adult offspring were collected for 3 days and then transferred to 29 °C to inactivate the temperature sensitive GAL80^ts^ protein. To generate MARCM or GFP-negative clones, flies were crossed at 25 °C and the resulting offspring were subjected to heat shocks either as larvae (from L2 until eclosion) or as adults (5–9 days after eclosion). Heat shocks were performed at 37 °C for 1 h twice daily. Flies were transferred to fresh food vials every 2–3 days and kept at 25 °C for at least 4 days after the last heat shock to ensure that all wild-type gene products from the heterozygous progenitor cells had turned over. All samples used in this study were obtained from adult female flies.

### Heat shock treatment in adult flies

Flies were subjected to heat shock in a 37°C incubator for 2hrs in a horizontal vial containing fly food but not live yeast. After removal from the incubator, live yeast was added to the vial and the flies were kept at 25C for 24hrs before dissection. To test the effect of heat shock treatment on ISC divisions, we counted the number of pH3^+^ cells in attp2 (Bloomington #25710) flies as control. Enteroblast nuclear volume was measured using *Su(H)>mCD8GFP* flies.

### Formaldehyde fixation and Heat Fixation

Samples were fixed as following the protocol of Chen et al. (2018). Samples were dissected in PBS and fixed with 8% formaldehyde (in PBS containing 0.1% Triton X-100) for 10 min at room temperature. Following several washes with PBS supplemented with 0.1% Triton X-100 (washing buffer), samples were incubated in PBS containing 3% normal goat serum (NGS, Stratech Scientific Ltd, Cat. #005-000-121; concentration of stock solution: 10 mg/ml) and 0.1% Triton X-100 (blocking buffer) for 30 min. This fixation method was only used for samples in which F-actin was stained with fluorescently labelled phalloidin, as phalloidin staining is incompatible with heat fixation. The heat fixation protocol is based on a heat– methanol fixation method used for *Drosophila* embryos (Muller, 2008). Samples were dissected in PBS, transferred to a wire mesh basket, and fixed in hot 1X TSS buffer (0.03% Triton X-100, 4 g/L NaCl; 95 °C) for 3 s before being transferred to ice-cold 1X TSS buffer and chilled for at least 1 min. Subsequently, samples were transferred to washing buffer and processed for immunofluorescence stainings.

### Antibody stainings

Antibody stainings were performed as described by Chen et al. (2018). After blocking in blocking buffer (1xPBS, 0.1%TritonX-100, 5%NGS (vol/vol)), samples were incubated with the appropriate primary antibody/antibodies diluted in blocking buffer at 4 °C overnight. Following several washes in washing buffer (1xPBS, 0.1%TritonX-100), samples were incubated with the appropriate secondary antibody/antibodies either at room temperature for 2 h or at 4 °C overnight. Samples were then washed several times in washing buffer and mounted in Vectashield containing DAPI (Vector Laboratories) on borosilicate glass slides (No. 1.5, VWR International). All antibodies used in this study were tested for specificity using clonal analysis (MARCM) or RNAi.

#### Primary antibodies

Mouse monoclonal antibodies: anti-Dlg (4F3), anti-Cora (c615.16), anti-αSpec (3A9), anti-Arm (N2 7A1), anti-Pros (MR1A). All monoclonal antibodies were obtained from the Developmental Studies Hybridoma Bank and used at 1:100 dilution. Rabbit polyclonal antibodies: anti-p4EBP1 (Phospho-4E-BP1 (Thr37/46) (236B4) Rabbit mAb #2855#2855T, Cell Signaling Technology), anti-pERK (Phospho-p44/42 MAPK (Erk1/2) (Thr202/Tyr204) Antibody #9101#9101s, Cell Signaling Technology), anti-β_H_-spectrin (gift from G. Thomas, Pennsylvania State University, USA, 1:1,000 dilution); anti-Par6 (gift from D. J. Montell, University of California Santa Barbara, USA, 1:500 dilution); anti-Mesh and anti-Tsp2A (gifts from Mikio Furuse, National Institute for Physiological Sciences, Okazaki, Japan, 1:1,000 dilution); anti-Pdm1 (gift from F. J. Diaz-Benjumea, Centre for Molecular Biology “Severo Ochoa” (CBMSO), Spain, 1:1,000 dilution); anti-Canoe (gift from M. Peifer, University of North Carolina, USA, 1:1,000 dilution); anti-Sox21a (Meng and Biteau, 2015)(gift from B. Biteau, University of Rochester, USA, 1:1000 dilution); anti-Rab11 (gift from Akira Nakamura, Kumamoto University, Japan) Other antibodies used: Chicken anti-GFP (Abcam, Cat. #ab13970, 1:1,000 dilution); Guinea pig anti-Myo7a (gift from D. Godt, University of Toronto, Canada, 1:1,000 dilution).

#### Secondary antibodies

Alexa Fluor secondary antibodies (Invitrogen) were used at a dilution of 1:1,000. Alexa Fluor 488 goat anti-mouse (#A11029), Alexa Fluor 488 goat anti-rabbit (#A11034), Alexa Fluor 488 goat anti-guinea pig (#A11073), Alexa Fluor 488 goat anti-chicken IgY (#A11039), Alexa Fluor 555 goat anti-mouse (#A21422), Alexa Fluor 555 goat anti-rabbit (#A21428), Alexa Fluor 568 goat anti-guinea pig (#A11075), Alexa Fluor 647 goat anti-mouse (#A21236), Alexa Fluor 647 goat anti-rabbit (#A21245). Only cross-adsorbed secondary antibodies were used in this study to eliminate the risk of cross-reactivity.

F-Actin was stained with phalloidin conjugated to Rhodamine (Invitrogen, Cat. #R415, 1:500 dilution).

### Immunofluorescence Imaging

Images were collected on an Olympus IX81 (40× 1.35 NA Oil UPlanSApo, 60× 1.35 NA Oil UPlanSApo) using the Olympus FluoView software Version 3.1 and processed with Fiji (ImageJ).

### Sample processing for electron microscopy and TEM imaging

Samples were fixed in fixative (2% glutaraldehyde/2% formaldehyde in 0.05M sodium cacodylate buffer pH7.4 containing 2mM calcium chloride) overnight at 4°C. After washing 5x with 0.05M sodium cacodylate buffer pH7.4, samples were osmicated (1% osmium tetroxide, 1.5% potassium ferricyanide, 0.05M sodium cacodylate buffer pH7.4) for 3 days at 4°C. After washing 5x in DIW (deionised water), samples were treated with 0.1% (w/v) thiocarbohydrazide/DIW for 20 minutes at room temperature in the dark. After washing 5x in DIW, samples were osmicated a second time for 1 hour at RT (2% osmium tetroxide/DIW). After washing 5x in DIW, samples were blockstained with uranyl acetate (2% uranyl acetate in 0.05M maleate buffer pH5.5) for 3 days at 4°C. Samples were washed 5x in DIW and then dehydrated in a graded series of ethanol (50%/70%/95%/100%/100% dry) 100% dry acetone and 100% dry acetonitrile, 3x in each for at least 5min. Samples were infiltrated with a 50/50 mixture of 100% dry acetonitrile/Quetol resin (without BDMA) overnight, followed by 3 days in 100% Quetol (without BDMA). Then, the sample was infiltrated for 5 days in 100% Quetol resin with BDMA, exchanging the resin each day. The Quetol resin mixture is: 12g Quetol 651, 15.7g NSA, 5.7g MNA and 0.5g BDMA (all from TAAB). Samples were placed in embedding moulds and cured at 60°C for 3 days.

Ultrathin sections (~70 nm) were cut on a Leica Ultracut E ultramicrotome and placed on bare 300 mesh copper TEM grids. Samples were viewed in a Tecnai G20 electron microscope (FEI/ThermoFisher SCientific) operated at 200 keV using a 20 μm objective aperture to enhance contrast. Images were acquired using an ORCA HR high resolution CCD camera (Advanced Microscopy Techniques Corp.).

### Generation of *FRT82B canoe*^JC^ flies

We used the CRISPR/Cas9 method [91] to generate a null allele of *canoe*. sgRNA was in vitro transcribed from a DNA template created by PCR from two partially complementary primers: forward primer: 5′-GAAATTAATACGACTCACTATA<u>GCAAACTCTCAGATGTGCAC</u>GTTTTAGAGCTAGAAATAGC-3′; reverse primer: 5′-AAAAGCACCGACTCGGTGCCACTTTTTCAAGTTGATAACGGACTAGCCTTATTTTAACTTGCTATTTCTAGCTCTAAAAC-3′. The sgRNA was injected into *Act5c-Cas9; FRT82B* embryos [92]. Putative *canoe* mutants in the progeny of the injected embryos were recovered, balanced, and sequenced. The *canoe*^JC1^ allele contains a small deletion around the CRISPR site, resulting in one missense mutation and a frameshift that leads to stop codon at amino acid 908 in the middle of the Dilute (DIL) domain, which is shared by all isoforms. No Canoe protein was detectable by antibody staining in both midgut and follicle cell clones.

### Generation of the *mesh*^R2^ flies

The *FRT82B cno*^R2^ flies were crossed with empty *FRT82* flies for homologous recombination. The progeny from this cross were balanced and complementation tests were performed by crossing with *FRT82B cno*^R10^ and *FRT82B mesh*^f04955^ to identify recombinant chromosomes that carried the *mesh* allele, but not *canoe*^R2^. Two homozygous lethal lines were obtained afterwards, named as *mesh*^R2^. Both mutant lines gave the same phenotype in clones in the adult midgut as *mesh*^f04955^.

### Measuring cortical Canoe intensities in *mesh*^R2^ mutant and wild-type cells

Fiji was used to measure cortical Canoe intensities in MARCM clones of *mesh*^R2^ and neighbouring wild-type integrating enteroblasts. We used the plot line tool to mark the cell contours (excluding the basal region), so that the centre value of the plot line represents the apical side and both peripheral values represent the basal side. Canoe intensities along the plot line are given as the percentage of the maximum Canoe intensity in the sample. The x-axis shows the position along the cell contour expressed as a percentage of the full-length value.

### Measuring Sox21a nuclear intensity in integrating enteroblasts in different stages

*sqh∷UtroABD-GFP* flies were heat shocked a day before dissection and heat-fixation. Rabbit anti-Sox21a was used to detect Sox21a nuclear signal. Sox21a nuclear intensity was measured at different stages of enteroblast integration relative to the apical actin signal using Fiji. The nuclear Sox21a level in enterocytes was measured in the same sample for comparison. The graph was made using GraphPad Prism 9 software.

### Counting the number of cells of various septate junction mutants in different stages of integration processes

SJ mutant cells (MARCM or negatively marked) were classified into five phenotypic classes: unpolarised cells, polarised cells, cells with a PAC, cells with an internal PAC and cells with an apical domain, based on their actin staining. The number mutant cells in each class was counted using Fiji and plotted using GraphPad Prism 9.

---

**Figure S1.**
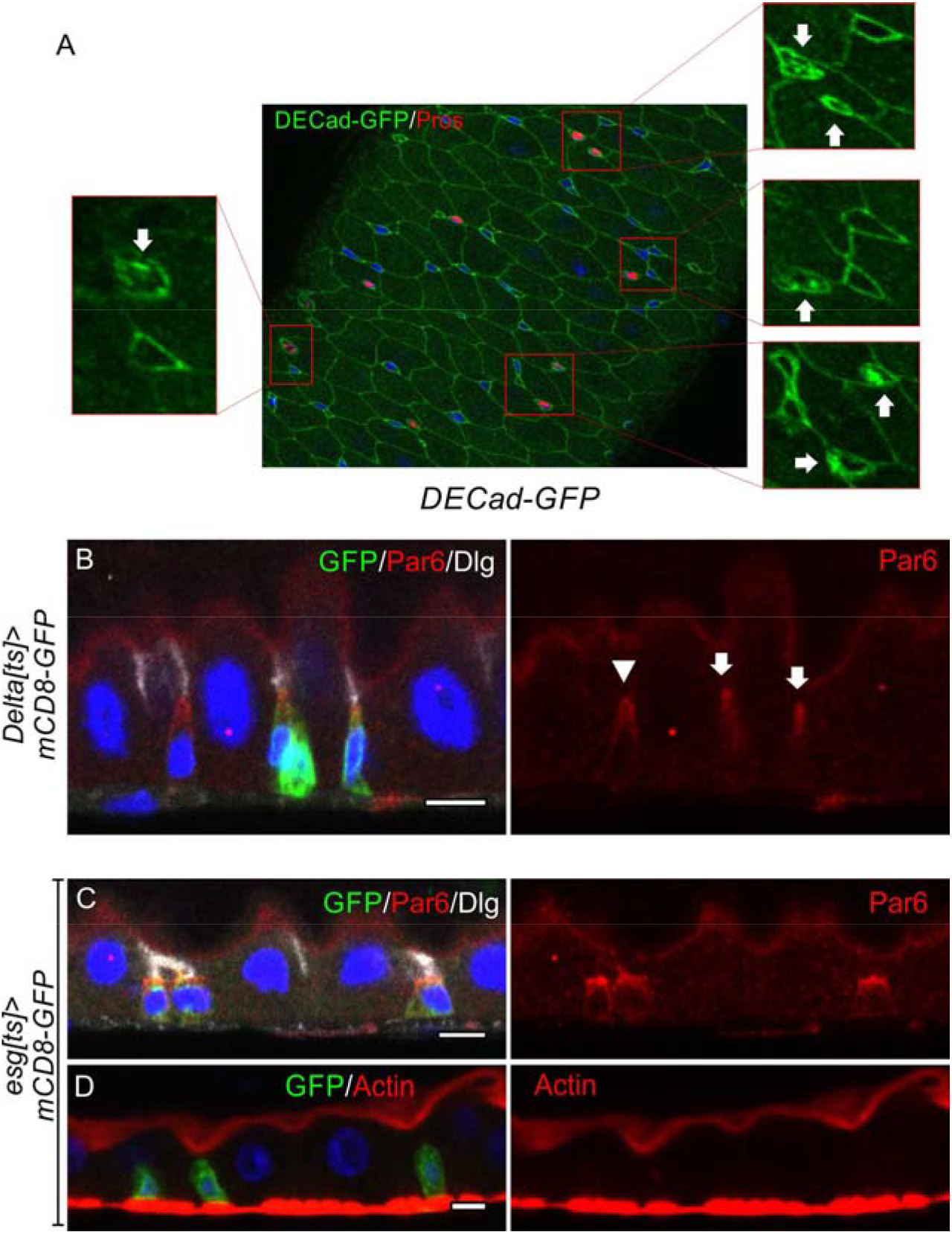
(related to Figure 1) (A) A midgut expressing DECadherin-GFP stained for GFP (green) and Prospero (red). The arrows indicate the Prospero^+^ entero-endocrine cells in the magnified images on each side. (B) Par-6 (red) is localised apically in ISCs (Delta^ts^>GFP positive cells; arrow) and early enteroblasts (Delta^ts^>GFP negative cell; arrowhead). (C-D) Par-6 (red; C) localises apically in esg^ts^>GFP positive ISC/early enteroblasts, whereas F-actin (red; D) is not apically-enriched. Dlg (greyscale; C) marks the septate junctions. Scale bars, 5μm.

**Figure S2.**
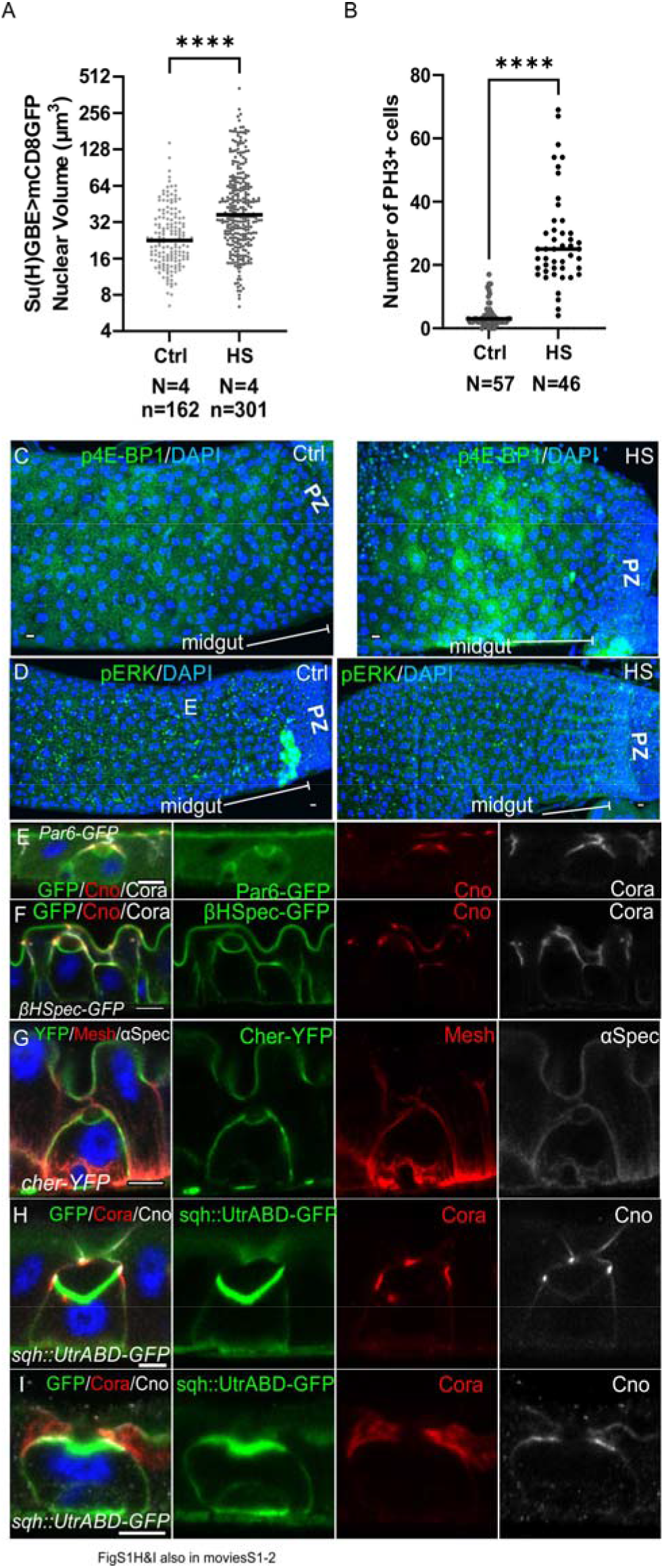
(related to Figure 2) (A) Graph showing the distribution of enteroblast nuclear volumes in control guts and in guts imaged one day after a 2 hour heat shock. The horizontal lines indicate the median values, which are significantly different by a 2-tailed t test (**** p< 0.0001). After heat-shock treatment, the Su(H)GBE>GFP+ enteroblasts have larger nuclear volumes on average, indicating that more cells are activated and becoming polyploid. N shows the number of midguts analysed in each experiment and n, the number of nuclei. (B) Graph showing the number of cells in mitosis (phospho-histone H3 (PH3) staining) in control guts and in guts imaged one day after a 2 hour heatshock. The heat shock increases the number of ISC divisions. The horizontal lines indicate the median values, which are significantly different by a 2-tailed t test (**** p< 0.0001). N shows the number of midguts analysed in this experiment. (C) Phosphorylated eukaryotic translation initiation factor 4E-binding protein 1 (p-4EBP1; green) staining in a control gut (left panel) and a gut 24 hours after a 2 hour heat shock (right panel). The heat shock induces a modest increase in p-4EBP1 levels. DNA is stained with DAPI (blue). (D) Phosphorylated Extracellular Regulated Kinase (p-ERK; green) staining in a control gut (left panel) and a gut 24 hours after a 2 hour heat shock (right panel). p-ERK levels are not increased after this treatment. (E) Par-6-GFP localisation to the lumen-facing domain of an integrating enteroblast. Canoe is labelled in (red) and the septate junctions are labelled with Coracle (greyscale). (F) β_H_-spectrin-GFP (green) localises to the enterocyte apical membrane and marks the lumen-facing membranes of both the integrating enteroblast and the overlying enterocytes. Canoe is labelled in (red) and the septate junctions are labelled with Coracle (greyscale). (G) A GFP protein trap line in the actin crosslinker, Cheerio (Cher; green). Cheerio localises to apical brush border of the enterocytes and to the apical, lumen-facing surface of an integrating enteroblast. Mesh (red) marks the septate junctions and basal labyrinth and α-spectrin (greyscale) marks the cell membranes. (H) Separate channels from the image of an integrating enteroblast with a closed pre-apical compartment (PAC) shown in Fig 2G. (I) Separate channels from the image of an integrating enteroblast with an open pre-apical compartment (PAC) shown in Fig 2H.

**Figure S3.**
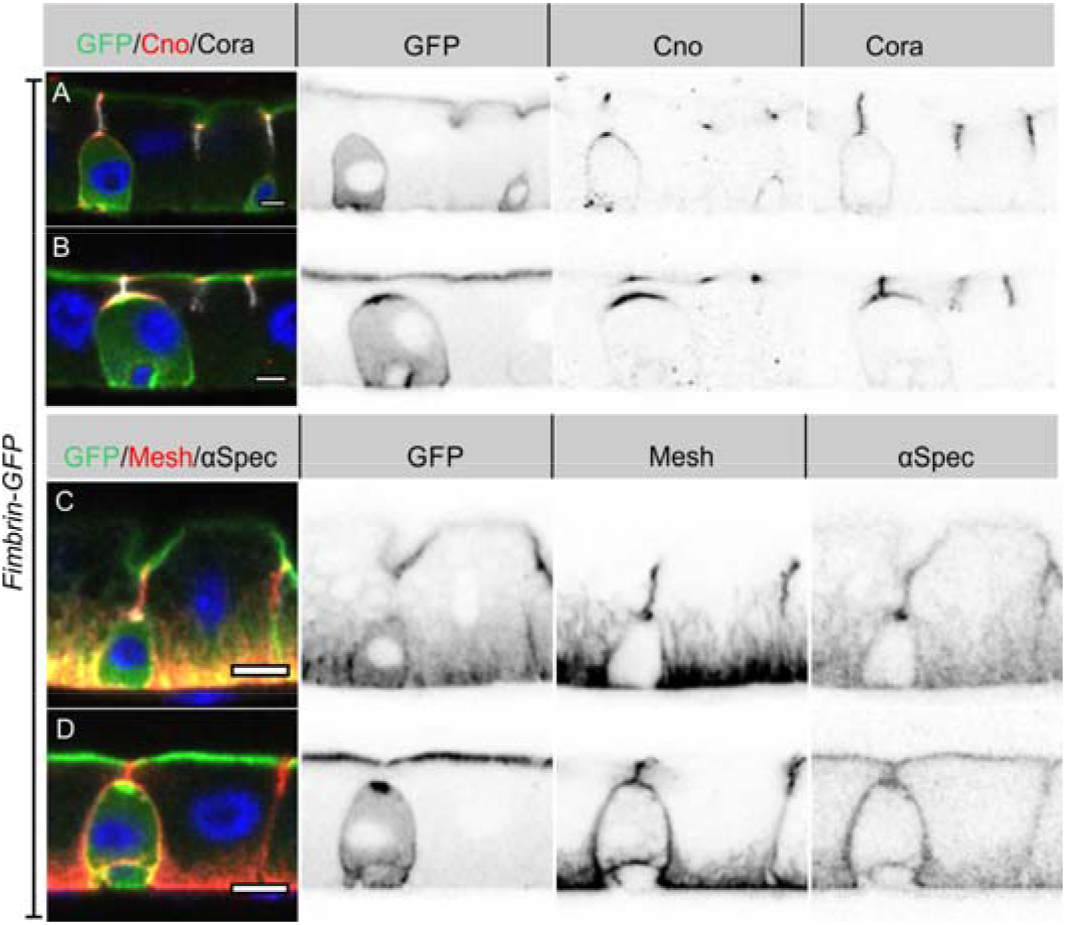
(related to Figure 3) (A-B) Fim-GFP (green) marking actin. The enteroblasts in A and B are at similar stages to those shown in in Fig 3A and 3C. Canoe (red) and Cora (greyscale). (C-D) Fim-GFP (green; actin), Mesh (red, septate junctions) and α-spectrin (greyscale; cell cortex). The enteroblasts in C and D are at similar stages as those in Fig 2A and 2C. Mesh also labels the membrane invaginations from the basal side that form the basal labyrinth. Scale bars = 5μm.

**Figure S4.**
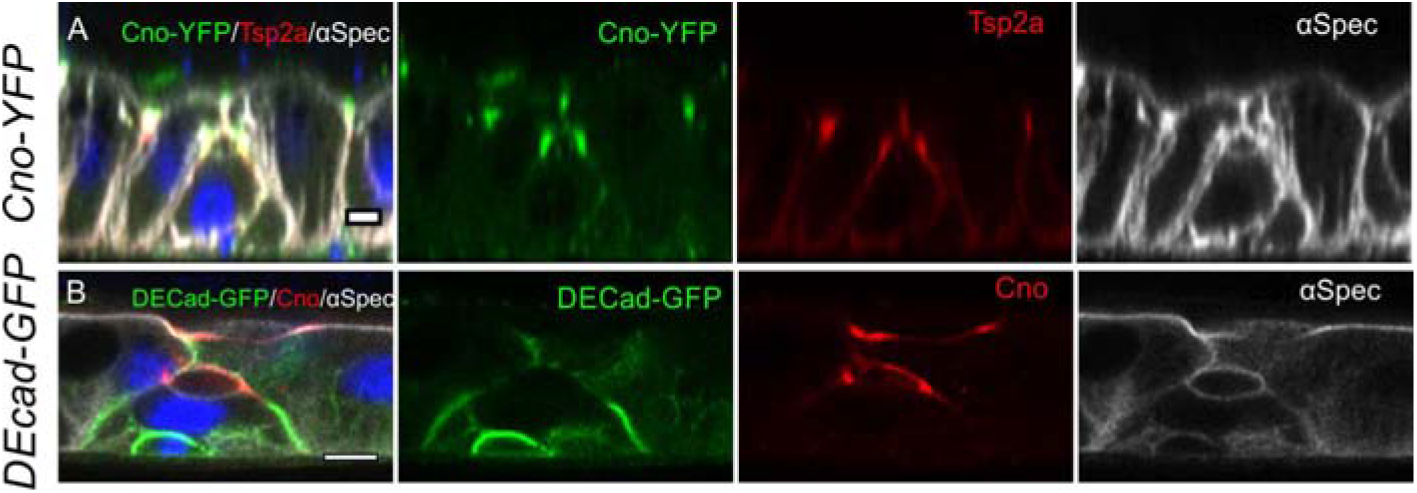
(related to Figure 4) (A) Tsp2a (red) localises to the septate junctions that develop between the pre-enterocyte and the neighbouring enterocytes as the PAC forms. Canoe is shown in green and α-spectrin in white. (B) E-cadherin (green) disappears around the PAC. Canoe is shown in red and α-spectrin in white. Scale bars, 5μm

**Figure S5.**
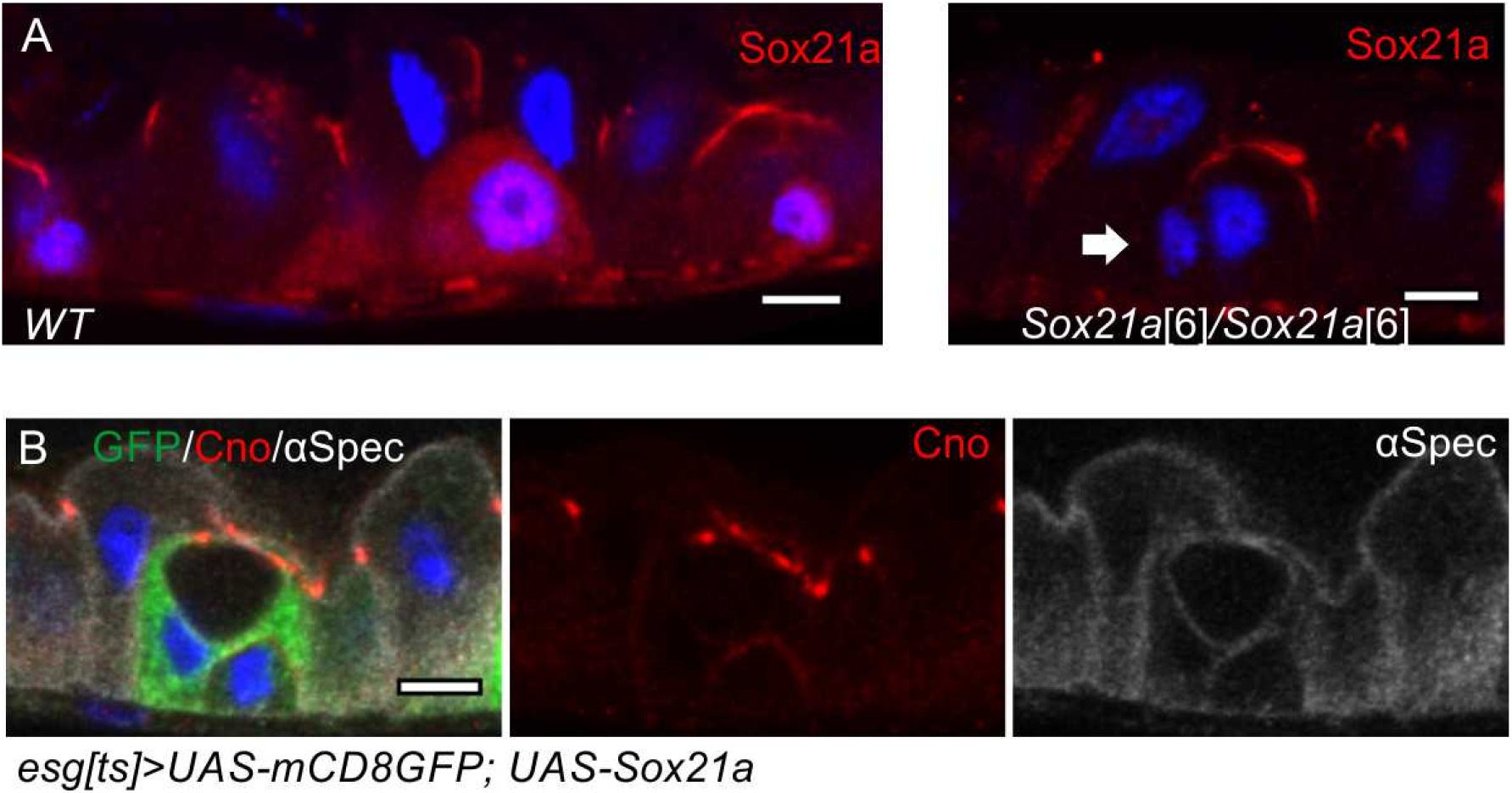
(related to Figure 5) (A) The Sox21a antiserum non-specifically stains the septate junctions. In the left hand panel, a wild-type Sqh∷UtrABD-GFP gut stained for Sox21a shows nuclear staining in the two enteroblasts, as well as strong staining of the septate junctions. The right hand panel shows a gut from *Sox21a*^6^ homozygote, in which the nuclear signal is lost, but the septate junctions are still labelled. The arrow points an enteroblast with no Sox21a nuclear signal. (A) An enteroblast over-expressing Sox21a that has formed a PAC, while remaining esg^ts^>GFP positive (green). Canoe in red and α-spectrin in greyscale. Scale bars = 5μm.

**Figure S6.**
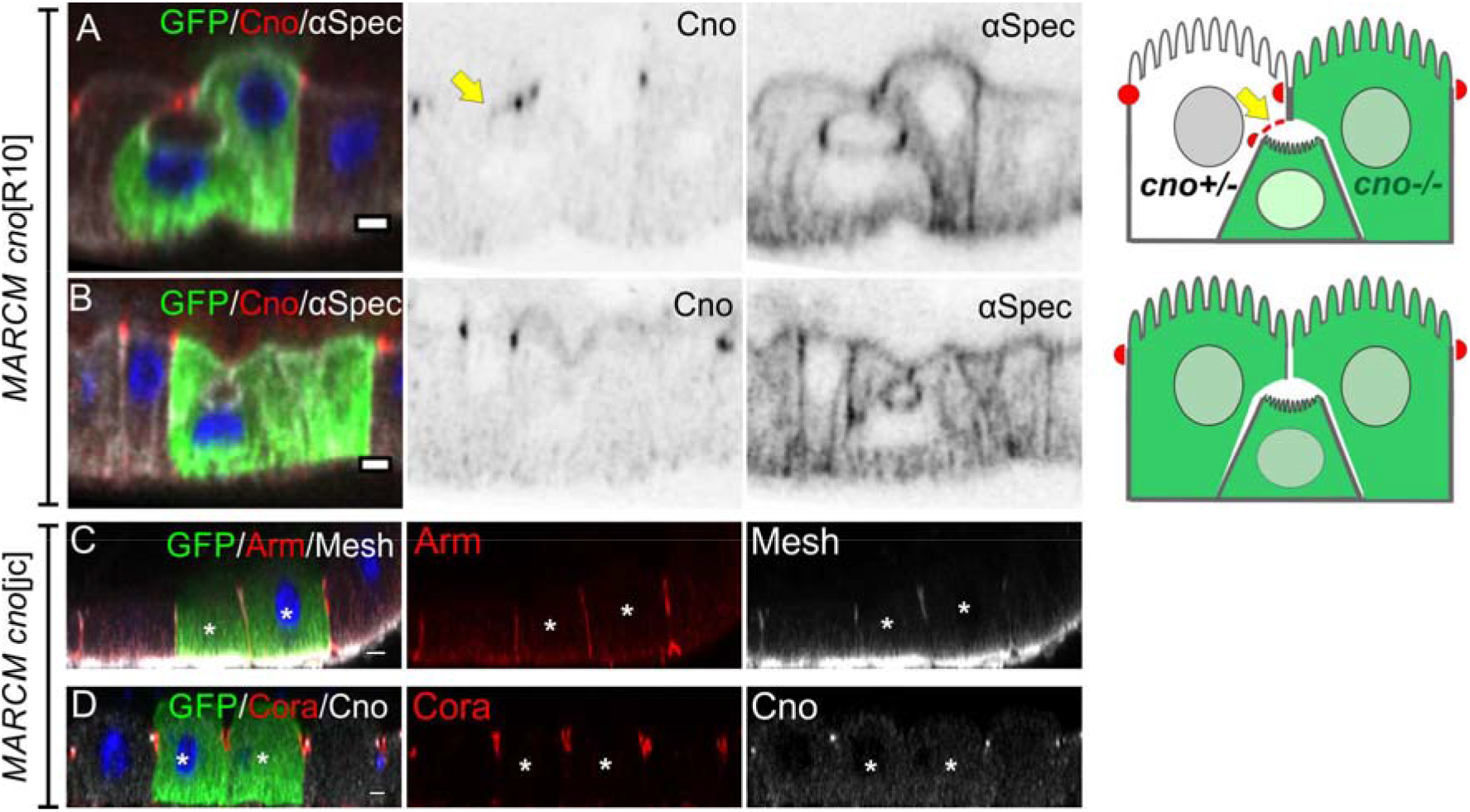
(related to Figure 7) (A) A PAC still forms when both the pre-enterocyte and one of the neighbouring enterocytes are homozygous for *canoe*^R10^ (green). Canoe in red and α-spectrin in white. (B) Canoe staining is lost completely in the PAC when all surrounding cells are *canoe*^R10^ mutant. (C-D) Adherens junctions (Arm, red in C) and septate junctions (Mesh, red in C; Cora, red in D) form normally in *canoe*^jc^ homozygous mutant cells (green). Scale bars, 5μm.

**Figure S7.**
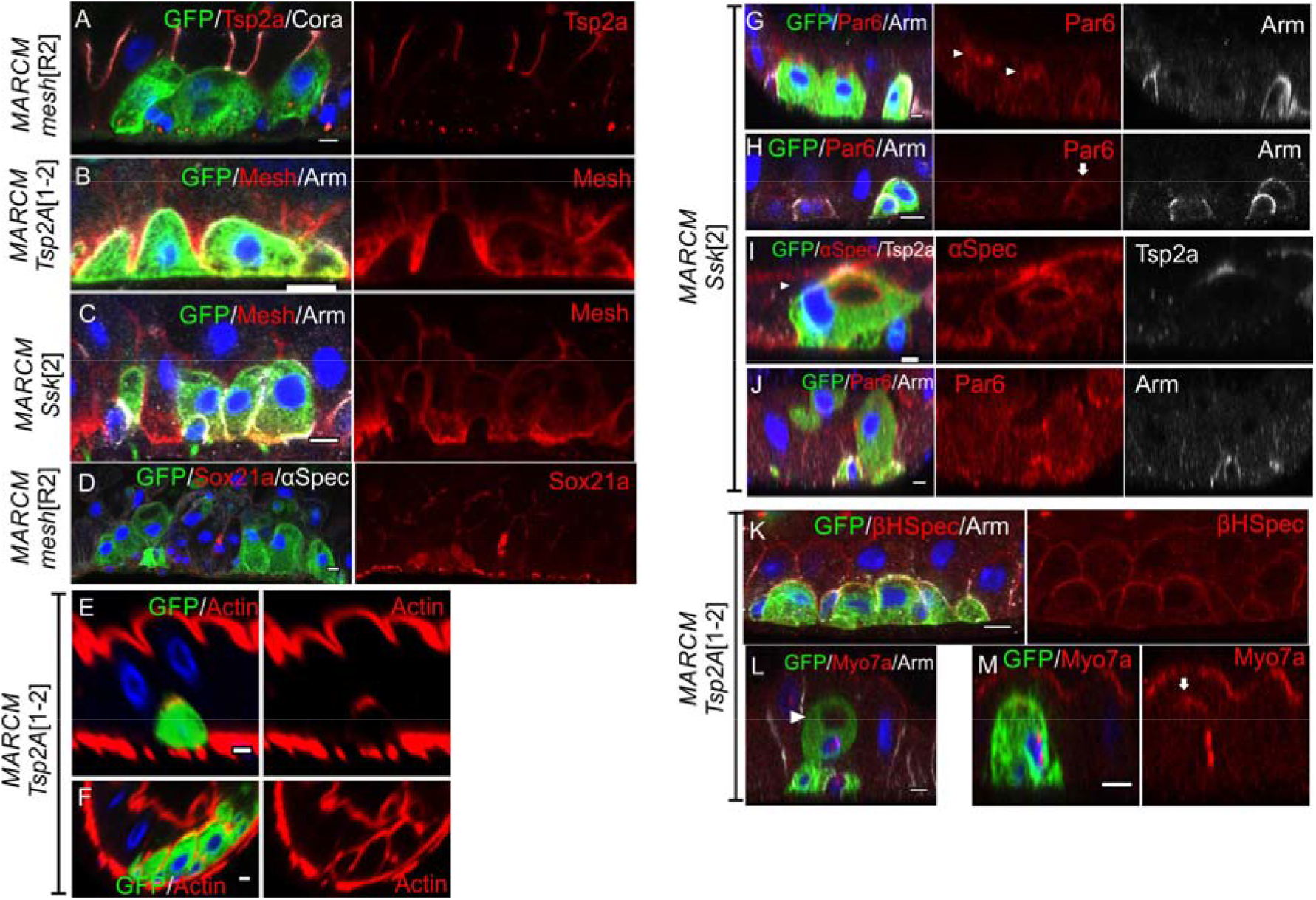
(related to Figure 8). (A-C) The localisations of septate junction proteins are interdependent. Tsp2A (red) is lost from *mesh*^R2^ mutant cells (green; A) and Mesh is not localised in *Tsp2A*^1-2^ (B) and s*sk*^2^ mutant cells (C). (D) Sox21a (red) is expressed and then down-regulated in *mesh*^R2^ mutant enteroblasts as in wild-type. (E-F) Representative images showing actin in two phenotypic classes of *Tsp2A*^1-2^ mutant cells as in Figure 8J: actin in polarised cells (E) and unpolarised cells (F). (G-J) *ssk*^2^ mutant cells still polarise (H), form a PAC (arrowhead in G) or an internal PAC (arrowhead in I) and occasionally reach the gut lumen (J). Par-6, red. (K-M) *Tsp2A*^1-2^ mutant enteroblasts (green) fall into three phenotypic classes: unpolarised (K), with an internal PAC (L) and weakly polarised (M). Scale bars, 5μm.

**Figure S8.**
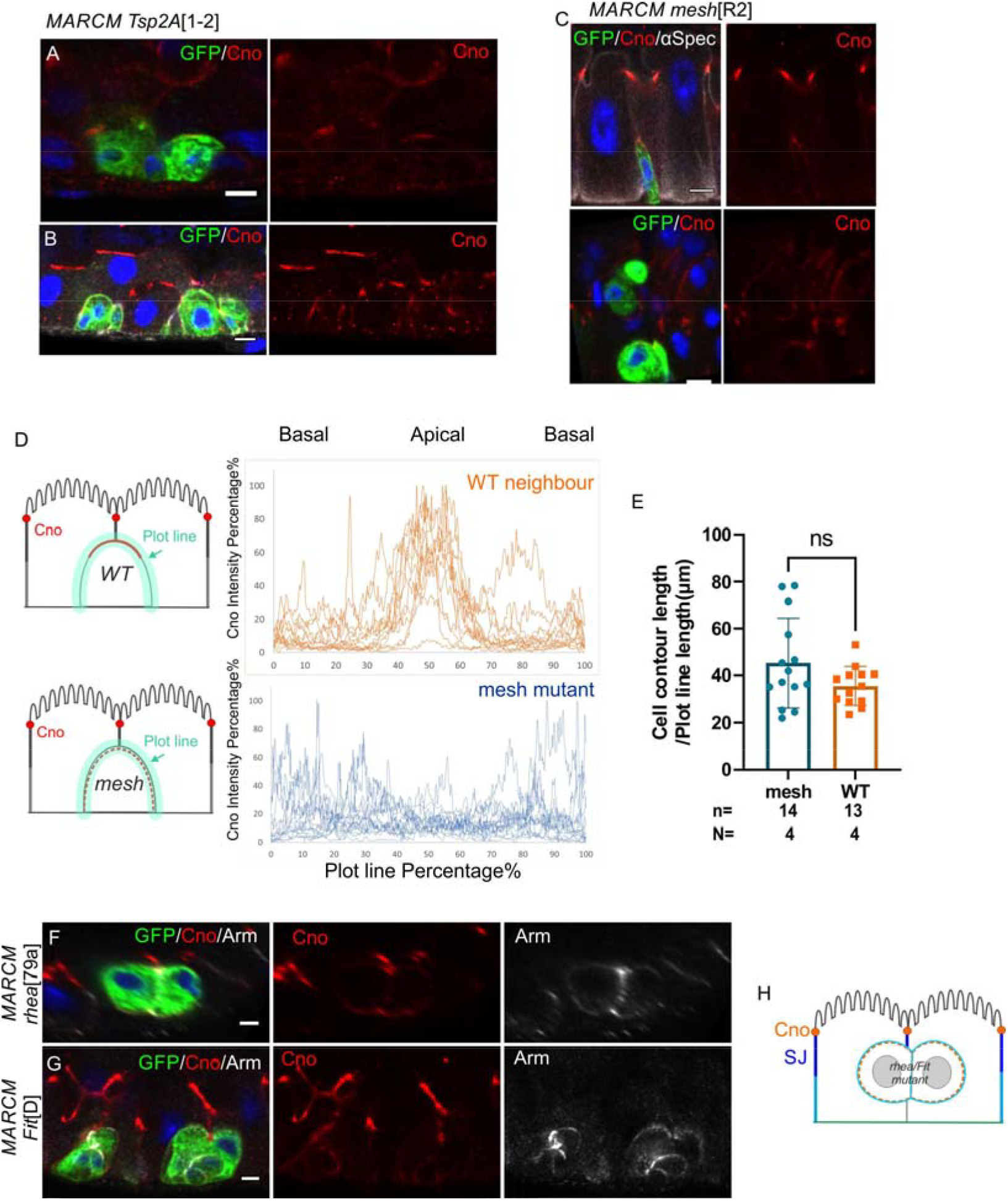
(related to Figure 8) (C) Canoe (red) is not polarised in larger *Tsp2A*^1-2^ mutant enteroblasts (green). (A-B) Canoe (red) is polarised in early stage *Tsp2A*^1-2^ (B) and *mesh*^R2^ mutant enteroblasts (C). (D) Graph showing the intensity of Canoe staining around the cell cortex from the basal (0% and 100%) to the apical side (50%) of 10 wild-type (brown) and 10 *mesh*^R2^ mutant cells. Canoe is enriched apically in wild-type cells, but not in *mesh*^R2^ mutant cells. (E) The measured *mesh*^R2^ and WT cells are similar in size. (F-G) Canoe (red) is not localised in *rhea*^79a^(F) and *Fit*^D^ (G) mutant cells. Scale bars = 5μm.

## Data availability

All fly stocks and reagents used in this study are available on request.

## Acknowledgements

We would like to thank Gurdon Institute Imaging Facility for microscopy and image analysis support, Karin Mueller and the Cambridge Advanced Imaging Centre for help with the transmission electron microscopy, Dr Jemima Burden (MRC_LMCB, UCL, London UK) for help with analysing TEM results, and members of the St Johnston laboratory for their advice and support. We are very grateful to Tony Ip, Mark Peifer, Thomas Lecuit, Stefan Luschnig, Benjamin Klapholz, Nicholas Brown, Golnar, Kolahgar, Mikio Furuse, Mihoko Tame, Graham Thomas, Denise Montell, Fernando Díaz-Benjumea, Benoit Biteau, Akira Nakamura, Dorothea Godt, the Bloomington *Drosophila* stock centre, the Kyoto Stock Center and the Developmental Studies Hybridoma Bank for providing fly stocks and antibodies.

This work was supported by a Wellcome Principal Fellowship (207496) to D. St J. and core funding from the Wellcome Trust (203144) and Cancer Research UK (A24843).

The authors declare no competing financial interests.

## Author contributions

J. Chen and D. St Johnston conceived and designed the project, prepared the figures and wrote and edited the manuscript. The experiments were performed by J. Chen. The project funding, administration, and supervision were provided by D. St Johnston.

## References

Amcheslavsky, A., Lindblad, J.L., and Bergmann, A. (2020). Transiently “undead” enterocytes mediate homeostatic tissue turnover in the adult *Drosophila* midgut. Cell Rep 33, 108408. 10.1016/j.celrep.2020.108408.

Antonello, Z.A., Reiff, T., Ballesta-Illan, E., and Dominguez, M. (2015). Robust intestinal homeostasis relies on cellular plasticity in enteroblasts mediated by miR-8-Escargot switch. EMBO J. 34, 2025–2041. 10.15252/embj.201591517.

Baumann, O. (2001). Posterior midgut epithelial cells differ in their organization of the membrane skeleton from other drosophila epithelia. Exp. Cell Res. 270, 176–187. 10.1006/excr.2001.5343.

Blasky, A.J., Mangan, A., and Prekeris, R. (2015). Polarized protein transport and lumen formation during epithelial tissue morphogenesis. Ann. Rev Cell Dev. Biol. 31, 575–591. 10.1146/annurev-cellbio-100814-125323.

Boettner, B., Harjes, P., Ishimaru, S., Heke, M., Fan, H.Q., Qin, Y., Van Aelst, L., and Gaul, U. (2003). The AF-6 homolog canoe acts as a Rap1 effector during dorsal closure of the Drosophila embryo. Genetics 165, 159–169. 10.1093/genetics/165.1.159.

Bryant, D.M., Datta, A., Rodríguez-Fraticelli, A.E., Peränen, J., Martín-Belmonte, F., and Mostov, K.E. (2010). A molecular network for de novo generation of the apical surface and lumen. Nat. Cell Biol. 12, 1035–1045. 10.1038/ncb2106.

Byri, S., Misra, T., Syed, Z.A., Bätz, T., Shah, J., Boril, L., Glashauser, J., Aegerter-Wilmsen, T., Matzat, T., Moussian, B., et al. (2015). The triple-repeat protein Anakonda controls epithelial tricellular junction formation in *Drosophila*. Dev. Cell 33, 535–548. 10.1016/j.devcel.2015.03.023.

Campbell, K., Whissell, G., Franch-Marro, X., Batlle, E., and Casanova, J. (2011). Specific GATA factors act as conserved inducers of an endodermal-EMT. Dev. Cell 21, 1051–1061. 10.1016/j.devcel.2011.10.005.

Chen, H.J., Li, Q., Nirala, N.K., and Ip, Y.T. (2020). The Snakeskin-Mesh complex of smooth septate junction restricts Yorkie to regulate intestinal homeostasis in *Drosophila*. Stem cell reports 14, 828–844. 10.1016/j.stemcr.2020.03.021.

Chen, J., and Sayadian, A.C. (2018). An alternative mode of epithelial polarity in the Drosophila midgut. PLoS Biology 16, e3000041. 10.1371/journal.pbio.3000041.

Chen, J., Xu, N., Huang, H., Cai, T., and Xi, R. (2016). A feedback amplification loop between stem cells and their progeny promotes tissue regeneration and tumorigenesis. 5. 10.7554/eLife.14330.

de Navascues, J., Perdigoto, C.N., Bian, Y., Schneider, M.H., Bardin, A.J., Martinez-Arias, A., and Simons, B.D. (2012). *Drosophila* midgut homeostasis involves neutral competition between symmetrically dividing intestinal stem cells. EMBO J. 31, 2473–2485. 10.1038/emboj.2012.106.

Engevik, A.C., Kaji, I., Postema, M.M., Faust, J.J., Meyer, A.R., Williams, J.A., Fitz, G.N., Tyska, M.J., Wilson, J.M., and Goldenring, J.R. (2019). Loss of myosin Vb promotes apical bulk endocytosis in neonatal enterocytes. J. Cell Biol. 218, 3647–3662. 10.1083/jcb.201902063.

Engevik, A.C., Krystofiak, E.S., Kaji, I., Meyer, A.R., Weis, V.G., Goldstein, A., Coutts, A.W., Melkamu, T., Saqui-Salces, M., and Goldenring, J.R. (2021). Recruitment of polarity complexes and tight junction proteins to the site of apical bulk endocytosis. Cell. Mol. Gast. Hep. 12, 59–80. 10.1016/j.jcmgh.2021.01.022.

Ferrari, A., Veligodskiy, A., Berge, U., Lucas, M.S., and Kroschewski, R. (2008). ROCK-mediated contractility, tight junctions and channels contribute to the conversion of a preapical patch into apical surface during isochoric lumen initiation. J. Cell Sci. 121, 3649–3663. 10.1242/jcs.018648.

Furuse, M., and Izumi, Y. (2017). Molecular dissection of smooth septate junctions: understanding their roles in arthropod physiology. Ann New York Acad. Sci 1397, 17–24. 10.1111/nyas.13366.

Goulas, S., Conder, R., and Knoblich, J.A. (2012). The Par complex and integrins direct asymmetric cell division in adult intestinal stem cells. Cell Stem Cell 11, 529–540. 10.1016/j.stem.2012.06.017.

He, L., Si, G., Huang, J., Samuel, A.D.T., and Perrimon, N. (2018). Mechanical regulation of stem-cell differentiation by the stretch-activated Piezo channel. Nature 555, 103–106. 10.1038/nature25744.

Izumi, Y., Furuse, K., and Furuse, M. (2019). Septate junctions regulate gut homeostasis through regulation of stem cell proliferation and enterocyte behavior in *Drosophila*. J. Cell Sci. 132. 10.1242/jcs.232108.

Izumi, Y., Furuse, K., and Furuse, M. (2021). The novel membrane protein Hoka regulates septate junction organization and stem cell homeostasis in the Drosophila gut. J. Cell Sci. 134. 10.1242/jcs.257022.

Izumi, Y., Motoishi, M., Furuse, K., and Furuse, M. (2016). A tetraspanin regulates septate junction formation in Drosophila midgut. J. Cell Sci. 129, 1155–1164. 10.1242/jcs.180448.

Izumi, Y., Yanagihashi, Y., and Furuse, M. (2012). A novel protein complex, Mesh-Ssk, is required for septate junction formation in the Drosophila midgut. J. Cell Sci. 125, 4923–4933. 10.1242/jcs.112243.

Jasper, H. (2020). Intestinal stem cell aging: origins and interventions. Ann. Rev. Physiol. 82, 203–226. 10.1146/annurev-physiol-021119-034359.

Jiang, H., and Edgar, B.A. (2011). Intestinal stem cells in the adult *Drosophila* midgut. Exp. Cell Res. 317, 2780–2788. 10.1016/j.yexcr.2011.07.020.

Jiang, H., Tian, A., and Jiang, J. (2016). Intestinal stem cell response to injury: lessons from *Drosophila*. Cell. Mol Life Sci. 73, 3337–3349. 10.1007/s00018-016-2235-9.

Klapholz, B., Herbert, S.L., Wellmann, J., Johnson, R., Parsons, M., and Brown, N.H. (2015). Alternative mechanisms for talin to mediate integrin function. Curr. Biol. 25, 847–857. 10.1016/j.cub.2015.01.043.

Lane, N.J., and Skaer, H.I. (1980). Intercellular junctions in insect tissues. Adv. Insect Physiol. 15, 35–213.

Lee, T., and Luo, L. (2001). Mosaic analysis with a repressible cell marker (MARCM) for Drosophila neural development. Trends in Neurosci. 24, 251–254. 10.1016/S0166-2236(00)01791-4.

Lowe, N., Rees, J.S., Roote, J., Ryder, E., Armean, I.M., Johnson, G., Drummond, E., Spriggs, H., Drummond, J., Magbanua, J.P., et al. (2014). Analysis of the expression patterns, subcellular localisations and interaction partners of *Drosophila* proteins using a pigP protein trap library. Development 141, 3994–4005. 10.1242/dev.111054.

Mangan, A.J., Sietsema, D.V., Li, D., Moore, J.K., Citi, S., and R., P. (2016). Cingulin and actin mediate midbody-dependent apical lumen formation during polarization of epithelial cells. Nat. Comms. 7, 12426. 10.1038/ncomms12426.

Meng, F.W., and Biteau, B. (2015). A Sox Transcription Factor Is a Critical Regulator of Adult Stem Cell Proliferation in the Drosophila Intestine. Cell Rep. 13, 906–914. 10.1016/j.celrep.2015.09.061.

Meng, F.W., Rojas Villa, S.E., and Biteau, B. (2020). Sox100B regulates progenitor-specific gene expression and cell differentiation in the adult *Drosophila* intestine. Stem Cell Rep 14, 226–240. 10.1016/j.stemcr.2020.01.003.

Micchelli, C.A., and Perrimon, N. (2006). Evidence that stem cells reside in the adult *Drosophila* midgut epithelium. Nature 439, 475–479. 10.1038/nature04371.

Miguel-Aliaga, I., Jasper, H., and Lemaitre, B. (2018). Anatomy and physiology of the digestive tract of *Drosophila melanogaster*. Genetics 210, 357–396. 10.1534/genetics.118.300224.

Muller, H.A. (2008). Immunolabeling of embryos. Meth. Mol. Biol. 420, 207–218. 10.1007/978-1-59745-583-1_12.

Müller, H.J. (2018). More diversity in epithelial cell polarity: A fruit flies’ gut feeling. PLoS Biology 16, e3000082. 10.1371/journal.pbio.3000082.

Nászai, M., Carroll, L.R., and Cordero, J.B. (2015). Intestinal stem cell proliferation and epithelial homeostasis in the adult *Drosophila* midgut. Insect Biochem Mol Biol. 67, 9–14. 10.1016/j.ibmb.2015.05.016.

O’Brien, L.E., Soliman, S.S., Li, X., and Bilder, D. (2011). Altered modes of stem cell division drive adaptive intestinal growth. Cell 147, 603–614. 10.1016/j.cell.2011.08.048.

Ohlstein, B., and Spradling, A. (2006). The adult *Drosophila* posterior midgut is maintained by pluripotent stem cells. Nature 439, 470–474. 10.1038/nature04333.

Ohlstein, B., and Spradling, A. (2007). Multipotent *Drosophila* intestinal stem cells specify daughter cell fates by differential Notch signaling. Science 315, 988–992. 10.1126/science.1136606.

Perez-Vale, K.Z., Yow, K.D., Johnson, R.I., Byrnes, A.E., Finegan, T.M., Slep, K.C., and Peifer, M. (2021). Multivalent interactions make adherens junction-cytoskeletal linkage robust during morphogenesis. J. Cell Biol. 220. 10.1083/jcb.202104087.

Pitsidianaki, I., Morgan, J., Adams, J., and Campbell, K. (2021). Mesenchymal-to-epithelial transitions require tissue-specific interactions with distinct laminins. J. Cell Biol. 220. 10.1083/jcb.202010154.

Reiff, T., and Antonello, Z.A. (2019). Notch and EGFR regulate apoptosis in progenitor cells to ensure gut homeostasis in Drosophila. EMBO J. 38, e101346. 10.15252/embj.2018101346.

Rojas Villa, S.E., Meng, F.W., and Biteau, B. (2019). zfh2 controls progenitor cell activation and differentiation in the adult Drosophila intestinal absorptive lineage. PLoS Genetics. 15, e1008553. 10.1371/journal.pgen.1008553.

Sasaki, A., and Nishimura, T. (2021). *white* regulates proliferative homeostasis of intestinal stem cells during ageing in *Drosophila*. Nat. Metab. 3, 546–557. 10.1038/s42255-021-00375-x.

Sawyer, J.K., Harris, N.J., Slep, K.C., Gaul, U., and Peifer, M. (2009). The Drosophila afadin homologue Canoe regulates linkage of the actin cytoskeleton to adherens junctions during apical constriction. J. Cell Biol. 186, 57–73. 10.1083/jcb.200904001.

Sedzinski, J., Hannezo, E., Tu, F., Biro, M., and Wallingford, J.B. (2016). Emergence of an Apical Epithelial Cell Surface In Vivo. Dev. Cell 36, 24–35. 10.1016/j.devcel.2015.12.013.

Shanbhag, S., and Tripathi, S. (2009). Epithelial ultrastructure and cellular mechanisms of acid and base transport in the Drosophila midgut. J. Exp. Biol. 212, 1731–1744. 10.1242/jeb.029306.

Tan, B., Yatim, S., Peng, S., Gunaratne, J., Hunziker, W., and Ludwig, A. (2020). The mammalian Crumbs complex defines a distinct polarity domain apical of epithelial tight junctions. Curr. Biol. 30, 2791–2804.e2796. 10.1016/j.cub.2020.05.032.

Taniguchi, K., Shao, Y., Townshend, R.F., Cortez, C.L., Harris, C.E., Meshinchi, S., Kalantry, S., Fu, J., O’Shea, K.S., and Gumucio, D.L. (2017). An apicosome initiates self-organizing morphogenesis of human pluripotent stem cells. J. Cell Biol. 216, 3981–3990. 10.1083/jcb.201704085.

Tepass, U. (2012). The apical polarity protein network in *Drosophila* epithelial cells: regulation of polarity, junctions, morphogenesis, cell growth, and survival. Ann. Rev. Cell Dev. Biol. 28, 655–685. 10.1146/annurev-cellbio-092910-154033.

Tepass, U., and Hartenstein, V. (1994). Epithelium formation in the *Drosophila* midgut depends on the interaction of endoderm and mesoderm. Development 120, 579–590.

Tepass, U., Tanentzapf, G., Ward, R., and Fehon, R. (2001). Epithelial cell polarity and cell junctions in *Drosophila*. Ann. Rev Gen. 35, 747–784. 10.1146/annurev.genet.35.102401.091415.

Wang, P., and Hou, S.X. (2010). Regulation of intestinal stem cells in mammals and Drosophila. J. Cell. Physiol. 222, 33–37. 10.1002/jcp.21928.

Wirtz-Peitz, F., Nishimura, T., and Knoblich, J.A. (2008). Linking cell cycle to asymmetric division: Aurora-A phosphorylates the Par complex to regulate Numb localization. Cell 135, 161–173. 10.1016/j.cell.2008.07.049.

Wu, K., Tang, Y., Zhang, Q., Zhuo, Z., Sheng, X., Huang, J., Ye, J., Li, X., Liu, Z., and Chen, H. (2021). Aging-related upregulation of the homeobox gene caudal represses intestinal stem cell differentiation in *Drosophila*. PLoS Genetics 17, e1009649. 10.1371/journal.pgen.1009649.

Xiang, J., Bandura, J., Zhang, P., Jin, Y., Reuter, H., and Edgar, B.A. (2017). EGFR-dependent TOR-independent endocycles support *Drosophila* gut epithelial regeneration. Nature Comms. 8, 15125. 10.1038/ncomms15125.

Xu, C., Tang, H.W., Hung, R.J., Hu, Y., Ni, X., Housden, B.E., and Perrimon, N. (2019). The Septate Junction Protein Tsp2A Restricts Intestinal Stem Cell Activity via Endocytic Regulation of aPKC and Hippo Signaling. Cell Rep. 26, 670–688.e676. 10.1016/j.celrep.2018.12.079.

Yarnitzky, T., and Volk, T. (1995). Laminin is required for heart, somatic muscles, and gut development in the Drosophila embryo. Dev. Biol. 169, 609–618. 10.1006/dbio.1995.1173.

Yu, H.H., and Zallen, J.A. (2020). Abl and Canoe/Afadin mediate mechanotransduction at tricellular junctions. Science 370. 10.1126/science.aba5528.

Zhai, Z., Boquete, J.P., and Lemaitre, B. (2017). A genetic framework controlling the differentiation of intestinal stem cells during regeneration in Drosophila. PLoS Genetics 13, e1006854. 10.1371/journal.pgen.1006854.

Zhai, Z., Kondo, S., Ha, N., Boquete, J.P., Brunner, M., Ueda, R., and Lemaitre, B. (2015). Accumulation of differentiating intestinal stem cell progenies drives tumorigenesis. Nature Comms. 6, 10219. 10.1038/ncomms10219.

